# An advanced head-to-tail mouse embryo model with hypoxia-mediated neural patterning

**DOI:** 10.1101/2025.06.17.660116

**Authors:** Anastasios Balaskas, Isabelle Kraus, Hatice Ö. Özgüldez, Persia Akbari Omgba, René Buschow, Adriano Bolondi, Idan Berlad, Jacob H. Hanna, Helene Kretzmer, Aydan Bulut-Karslioğlu

## Abstract

The developing mammalian embryo is guided by the continuously changing signals that it receives from maternal tissues and its microenvironment. The dynamic cell-cell and cell-environment interactions that together shape the embryo largely remained unexplorable until the advance of stem cell-based embryo models. These revealed the self-organizing properties of cells in response to endogenous and exogenous cues. Among the latter, restricted oxygen (hypoxia) emerged as a critical microenvironmental regulator that influences cell type diversification in multicellular systems. Here we built a modular ESC-based head-to-tail model of mouse embryogenesis by developing an <u>a</u>ntero-<u>p</u>osterior (AP) assembly strategy under hypoxia. These structures called HAP-gastruloids feature stage-appropriate anterior neural tissues that recapitulate the morphological organization and transcriptional identity of fore- and midbrain including spatial organizer regions such as the midbrain-hindbrain boundary. These anterior tissues develop in synchrony with posterior tissues such as the spinal cord, somites, and gut endoderm derivatives, ultimately yielding a unified structure. We show via genetic, environmental, and pharmacological perturbations that timed hypoxia is essential to boost anterior neural cell identities and their patterning through HIF1a and in part by modulating TGFβ activity. These results underline the key beneficial role of hypoxia in early development and offer a uniquely modular system to investigate antero-posterior phenotypes for basic discovery and translation.

## INTRODUCTION

The uterine habitat of the mammalian embryo nourishes, shapes, and guides the developing cells. Gas and nutrient exchange through the placenta is a vital part of mammalian development, yet, the earliest and most elemental stages of development take place before the placenta is formed and becomes fully functional. One such stage is gastrulation, during which the mammalian body plan is laid out, and the founding cells of all tissues are specified and initially localized. Over four decades of developmental biology research identified key signaling events that lay out the mammalian body plan^1,2^, yet, the investigation of how the uterine microenvironment contributes to these complex and concurrent events is an emergent question. Specifically, how the presumably low-oxygen (hypoxic) environment resulting from the lack of developed vasculature and a functioning placenta impacts the complexity of the gastrulating embryo is poorly understood^3–5^.

In the recent decade, various 2D/3D stem cell-based in vitro models of the developing embryo have been developed. The 3D models in particular revealed the remarkable self-organizing events in cellular aggregates, often yielding structures that closely recapitulate embryonic events under minimal guidance. The extent to which these structures present complex embryonic tissues along the three axes of symmetry depends largely on the complexity of the initial group of cells. The so-called ‘integrated’ models are generated using embryonic and extraembryonic stem cells and model the entire mammalian conceptus, including extra-embryonic tissues^6^, while ‘non-integrated’ models, generated using embryonic stem cells (ESCs), robustly recapitulate specific aspects of embryonic development^7–25^. Among the latter, gastruloids represent posterior gastrulation events and have been widely adopted to investigate cellular decision-making and self-organization properties^14^. However, a major limitation of gastruloids and gastruloid-derivatives models^10,13,26,27^ is the lack of brain specification and insufficient representation of other anterior cell fates. An effort has been made to generate anterior tissues in gastruloids^28–30^, however, a gap persists for the generation of sufficiently developed and morphologically organized structures with an anterior brain-like domain.

We have previously shown that the combined exposure of gastruloids to 2% oxygen (O_2_) and the WNT activator Chiron increases the cellular complexity of conventional gastruloids, whereas exposure to 2% O_2_ in the absence of Chiron allows the generation of anterior tissues with transcriptional signatures of the brain^31^. Additionally, timed and gradual exposure of embryos to low oxygen concentrations during *ex utero* mouse gastrulation results in higher rates of normally developed embryos compared to normoxia^32^. Deletion of the hypoxia-inducible factor 1a gene (*Hif1a*) in mice results in gestational deficiencies that go beyond the expected vascularization defects and extend into neural tube and forebrain formation^33^. Furthermore, within the rat neural lineage, different types of neurons can be generated under differing O_2_ concentrations^3,34,35^. Together, oxygen appears to modulate cell fates in a concentration-dependent manner comparable to a developmental morphogen.

Leveraging these effects of oxygen concentration in the context of embryonic assembloids, here we developed hypoxic A-P (HAP) gastruloids featuring anterior brain-like tissues with stage-appropriate regional patterning. HAP-gastruloids present a robust and modular embryo model that illuminates the early organogenesis stages.

## RESULTS

### HAP-gastruloids form all three germ layer derivatives including a brain-like domain

To test the limits of cell type diversification via hypoxia in a self-organizing early embryonic differentiation system, we first employed the gastruloid-like embryo model called Trunk Like Structures (TLS)^10,13^. TLSs are formed from a single, transgene-free ESC aggregate over 120 hours. A transient pulse of the WNT agonist Chiron between 48-72h allows symmetry breaking and axial elongation. Addition of extracellular matrix in the form of matrigel in the last 24h enables the emergence of stage-appropriate morphological features of the embryo posterior such as the neural tube, somites, and occasionally, a gut tube (Figure S1A-D). TLSs, however, entirely lack more anterior tissues such as the brain^36,37^. To test the boundaries of hypoxia-driven cell fate commitment and morphogenesis especially towards anterior neural tissues of the brain, we systematically tested the effect of hypoxia by altering its duration and severity before and during TLS formation (Figure S1A-D, conditions I-IX). We found that, early hypoxia exposure, followed by a gradual transition to normoxia enriched the overall cell type composition and morphology and was superior to the other tested conditions (Figure S1A-D, section ‘Optimization of O_2_ implementation in TLS model’ in *Methods* for details). However, we also concluded that starting with a single aggregate in this system is incompatible with recapitulating the entire A-P axis of the corresponding stage embryo. Generating well-developed tissues anterior to the somites appeared particularly challenging. The gastrulating mouse embryo creates a WNT gradient with high WNT activity at the posterior end, which is antagonized at the anterior end, allowing the generation of anterior neural cells that are precursors of the future brain. Reasoning that an analogous gradient *in vitro* may allow the formation of stage-appropriate cells along the A-P axis, we designed a new, assembly-based experimental approach.

Assembly-based models (assembloids) successfully recapitulate the morphological and compositional complexity of other mammalian tissues, such as the cortical-motor and cortical-vascular tissues of the brain^38–45^. To generate a model with an inherent WNT gradient along the A-P axis, we assembled a Chiron-treated aggregate with a Chiron-free aggregate in 1:1 ratio (conditions X and XI, Figure S2A). The two aggregates were placed in the same well at 72h, were completely fused by 96h into a single structure, which then elongated in the next 24h. All cells in this system experience the same oxygen gradient during differentiation. Notably, A-P assembloids robustly formed FOXC1^+^ somites, a SOX2^+^ posterior neural tube as well as an anterior neural domain positive for the midbrain/forebrain marker OTX2 (Figure S2B-C). These results support the A-P assembly approach as a means to modularly build the embryonic axis.

Further efforts to enhance tissue complexity under hypoxia revealed that the continuity of the neural tube is incompatible with robust formation of gut endoderm in this 2-aggregate system (Figure S2B, conditions X and XI). A 3-aggregate A-P assembly model partially overcame this limitation and simultaneously formed a continuous neural tube (70% of structures), an anterior brain-like domain (85%) along with gut endoderm derivatives (37% cavities, 48% individual gut tube rosettes) (Figure S2A-G, condition XII). A further iteration of adjusting cell numbers was done to equalize the size of each aggregate (Figure S2D). The resulting embryo model, which we term <u>h</u>ypoxic <u>A-P gastruloid</u> (**HAP-gastruloid**), contains somites, gut precursors and a continuous neural tube with spinal cord features on the posterior side and brain-like expression profile on the anterior side (Figure 1A-D). These features were reproduced using two independent ESC lines (Figure 1B-D, S3A-B).

**Figure 1:**
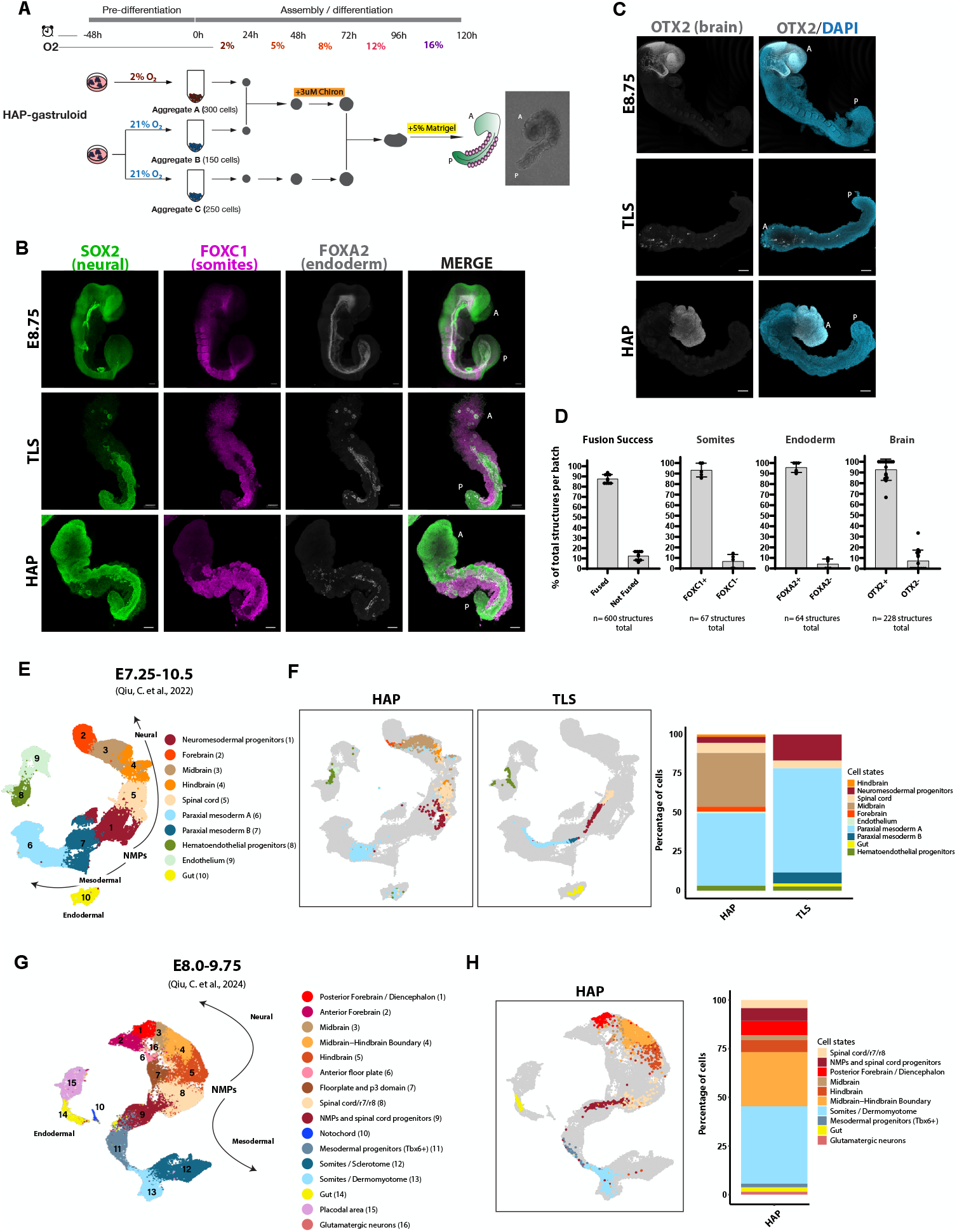
HAP-gastruloids form all three germ layer derivatives and a brain-like domain. A. Schematic illustration of the experimental pipeline for HAP-gastruloid generation. B. Immunofluorescence (IF) staining of E8.75 mouse embryo, TLS and HAP-gastruloids for neural marker SOX2, somitic marker FOXC1 and endoderm marker FOXA2. A: Anterior, P: Posterior. Scale bars, 100 μm. C. IF staining of mouse embryo at E8.75, TLS and HAP-gastruloids for brain marker OTX2 and DAPI. A: Anterior, P: Posterior. Scale bars, 100 μm D. Percentage of total structures per biological replicate showing fusion success, somite, endoderm and brain formation in HAP-gastruloids. Column heights show mean, and error bars indicate standard deviation. Each dot represents a different batch. E. UMAP plot of mouse embryo reference scRNA-seq data for stages between E7.25 to E10.546. Cell types that are robustly detected in HAP-gastruloids and/or TLSs are shown (>25 cells). Arrows show lineage trajectories. NMP, neuromesodermal progenitors. F. Left, UMAP plot for HAP-gastruloid and TLS cells projected onto the embryo atlas. Right, stacked bar plots showing cell type compositions. G. UMAP plot of mouse embryo reference scRNA-seq data for stages between E8.0 to E9.7547. Cell types that are detected in HAP-gastruloids are shown. Arrows show lineage trajectories. H. Left, UMAP plot of HAP-gastruloids projected onto the embryo atlas. Colors represent cell types. Right, stacked bar plots showing cell type compositions.

To probe gene expression patterns and cellular composition in more detail, we next performed single cell RNA-sequencing (scRNA-seq) on HAP-gastruloids and compared it to TLS. 6-20 structures per condition were pooled, yielding 3391 HAP-gastruloid cells and 2710 TLS cells after quality control. To gauge the developmental progress and cell type composition of HAPs, we first used an expanded scRNA-seq atlas of the embryonic stages E7.25-E-10.5 (Figure 1E-F, S4A)^46^. These analyses validated our stainings and suggest that, in contrast to TLS, HAP-gastruloids form brain-like tissues with transcriptional signatures of forebrain, midbrain and hindbrain. HAPs also feature the posterior tissues formed in canonical gastruloids/TLSs such as neuro-mesodermal progenitors (NMPs), somitic paraxial mesoderm tissue, spinal cord (neural tube), and gut endoderm. As a result, the proportion of posterior and anterior tissues is more balanced in HAP-gastruloids (Figure 1F, right). The transcriptional signatures of these tissues match the E8.5 embryo according to this atlas (Figure S4B).

To characterize the tissues in HAP-gastruloids better and cross-check these findings, we used a second scRNA-seq atlas which better resolves the developmental time window around E8.5 with embryonic samples collected at a 0.25-day interval between E8.0-E9.75 (Figure 1G-H, see S4C for gene markers of specific clusters)^47^. This atlas confirmed the notably enriched and balanced tissue representation in HAPs. The atlases show differences in specific annotations of brain-like HAP cells, with particularly the midbrain and hindbrain cells of the first atlas assigned to the midbrain/hindbrain boundary in the second atlas. Additionally, paraxial mesoderm was annotated as dermomyotome (dorsal fate) and mesodermal progenitors. In conclusion, based on both atlases HAP-gastruloids contain and spatially organize an expanded range of embryonic tissues along the entire A-P axis, revealing a capacity for anterior-posterior cross-talk and self-organization that results in a unified embryo-like structure starting from ESCs.

### HAP-gastruloid aggregates develop synchronously and compose diverse tissues

Since HAP-gastruloids are generated from the fusion of three different aggregates, a major outstanding question is whether the developmental timing of the distinct aggregates are synchronized to each other. For this, we first aimed to find out the specific contribution of each aggregate to the overall system. To achieve this, we used a SOX2::Venus and T::mCherry dual reporter ESC line^10^, and formed HAP-gastruloids by labeling one aggregate at a time (Figure S5A). This cell line labels cells with current or recent SOX2 expression in green and T (Brachyury) expression in red. We observed that the Chiron-treated aggregates (A and B, conditions I and II) both contributed to a range of posterior tissues. Among these, aggregate A mainly contributed to tissues surrounding the posterior neural tube and somites, whereas aggregate B formed the posterior neural tube (spinal cord) and somites themselves. The Chiron-free aggregate C exclusively formed the brain-like anterior region and also contributed to anterior spinal cord. Occasionally, this contribution even extended to more posterior neural regions (Figure S5A, condition III). Thus, although the brain-like anterior end was generated only by aggregate C, interactions of cells originating from all three aggregates could be observed along a large portion of the A-P axis.

The three aggregates develop independently until the moment of assembly. The overall developmental timing of the unified HAP-gastruloid matches the E8.5 embryo per transcriptional analyses of both atlases (Figure S5B, C). To next address whether the diverse tissues within HAP-gastruloids are synchronized with each other, we performed a label transfer-based staging analysis^48^ and plotted the predicted stage of each cell in each tissue for both references (Figure S5D). According to the first reference^46^, all cell types progressed synchronously to E8.5 transcriptionally, with a slightly later predicted timing only in the forebrain. In the second reference^47^, the predicted stages are more variable, ranging between E8.5-8.75 for most tissues and E8.75-E9.25 for posterior forebrain and midbrain. These variations were not observed in the respective midbrain and forebrain cells of the first reference. Thus, both cell type annotation and developmental stage prediction is prone to variability across atlases. Based on this comprehensive analysis, we conclude that HAP-gastruloids contain stage-appropriate cell types along the A-P axis, which together form a substantially developed embryo model of ESC origin that recapitulates aspects of E8.5-E8.75 embryonic development.

### HAP-gastruloids show regional neural patterning along the A-P axis

The most unique feature of HAP-gastruloids is the presence of a brain-like domain anterior to a continuous neural tube. More specifically, scRNA-seq analysis suggested the presence of specific regions within the brain-like domain, such as the posterior forebrain, midbrain and midbrain-hindbrain boundary (Figure 1G-H). The E8.5-9.0 mouse embryo contains primordial forebrain and midbrain regions that express PAX6 and OTX2 or only OTX2, respectively (Figure 2A). Posterior to these, PAX6 is also expressed in the spinal cord, while GBX2 is expressed in the anterior spinal cord (future hindbrain) and the tail. To test whether HAP-gastruloids indeed pattern the neural cells along the A-P axis, we stained HAP-gastruloids for PAX6, OTX2, and GBX2 (Figure 2B, S6A-B). Similar to the embryo, we find a PAX6+ domain within the OTX2+ region, indicating the specification of forebrain and midbrain regions (Figure 2B, C). Within the developing brain, molecular boundaries mark and define the regionalization of compartments. Among these, the FGF8, EN1, and WNT1 define the midbrain/hindbrain boundary. Staining for FGF8 and EN1 in HAP-gastruloids revealed a sharp boundary set by FGF8, with appropriate expression of EN1 around it (Figure 2D, E). This spatial patterning is a feature of the assembled HAP-gastruloid and is not observed in the stand-alone ‘brain’ aggregate (Figure 2F), indicating that the fusion and subsequent interaction of the posterior and anterior aggregates is essential for this regional neural patterning. These results were confirmed in two independent ESC lines (Figure S6A-D).

**Figure 2:**
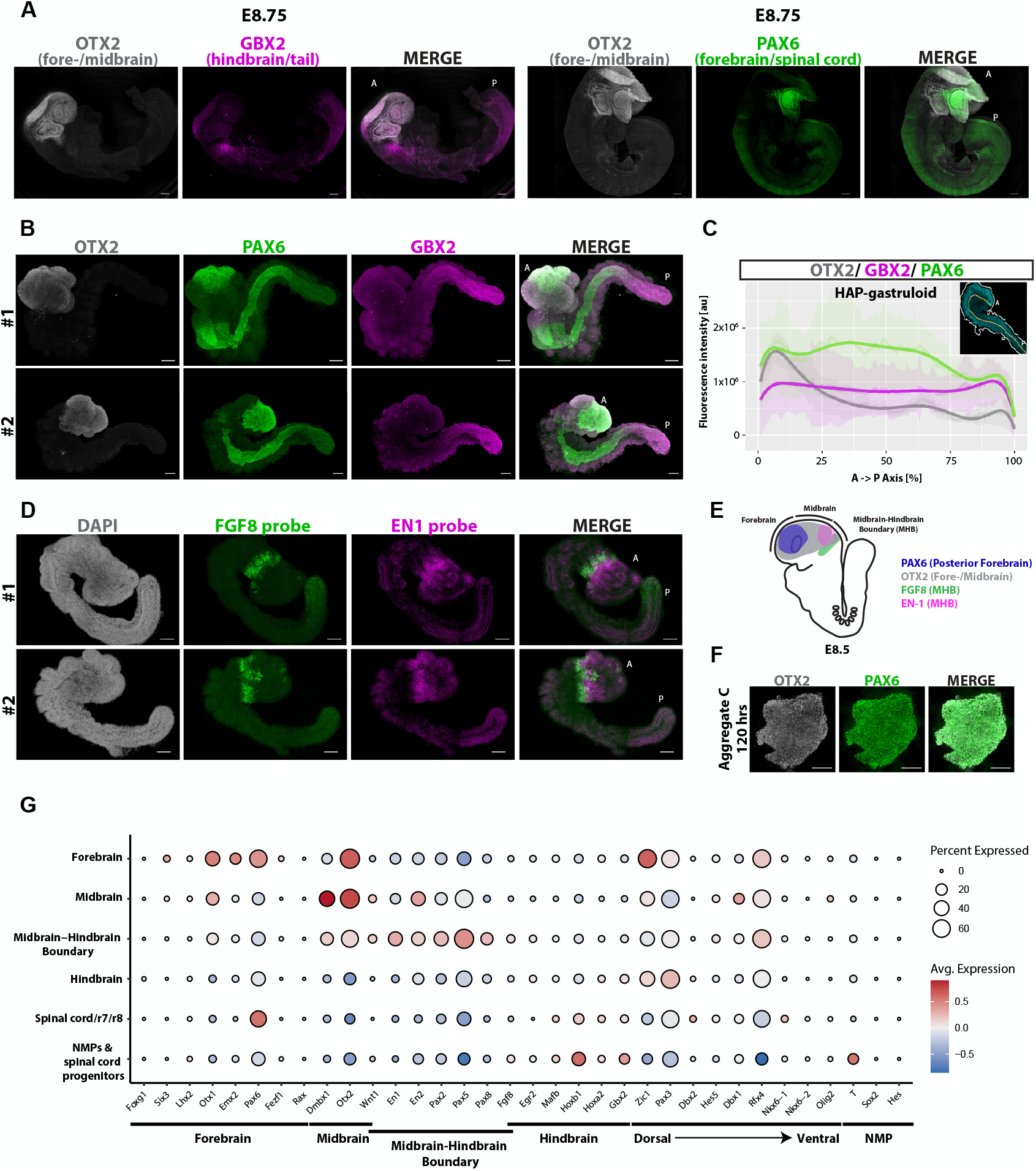
HAP-gastruloids show regional neural patterning along the A-P axis. A. IF staining of E8.75 mouse embryos for the fore-/midbrain marker OTX2, hindbrain/NMP marker GBX2 and forebrain/spinal cord marker PAX6. A: Anterior, P: Posterior. Scale bars, 100 μm. B. IF staining of HAP-gastruloids for fore-/midbrain marker OTX2, hindbrain/NMP marker GBX2 and forebrain/spinal cord marker PAX6. Scale bars, 100 μm. C. Localization of OTX2, GBX2 and PAX6 signals along the A-P axis of HAP-gastruloids. Lines show mean values from 9-11 individual structures. Small figure: line shows how the A-P axis was defined in structures. D. HCR RNA FISH of HAP-gastruloids for midbrain-hindbrain boundary markers EN1 and FGF8. Scale bars, 100 μm. E. Schematic Illustration of forebrain marker PAX6, fore-/midbrain marker OTX2 and midbrain-hindbrain boundary markers EN1 and FGF8 expression patterning along the A-P axis in E8.5 mouse brain. F. IF staining of aggregate C (Chironless) at 120 hrs for fore-/midbrain marker OTX2 and forebrain/spinal cord marker PAX6. Scale bars, 100 μm. G. Dot plot showing the expression of neural cell type specific marker genes in HAP-gastruloids47. Dot size corresponds to the percentage of cells expressing the gene, color intensity reflects the scaled average expression level.

To analyze the anterior neural tissues of HAP-gastruloids in more detail, we investigated marker expression of each brain compartment in the scRNA-seq data (Figure 2G). According to this, HAP-gastruloids express midbrain markers Dmbx1 and Wnt1 as well as midbrain-hindbrain boundary markers Pax2, Pax5, Pax8 and En2. Posterior forebrain markers Pax6 and Emx2 are expressed, but not markers of the most rostral forebrain such as Foxg1, Six3 and Lhx2. Hindbrain markers Gbx2 and Egr2 show mild expression. Ventral neural tube markers, Nkx6.1, Nkx6.1 and Olig2 are absent and thus HAP-gastruloids recapitulate mainly dorsal patterning along the A-P axis. Together, the combined hypoxia-assembloid strategy yields a regional dorsal neural patterning along the A-P axis, which is not seen in TLSs or stand-alone neural aggregates.

Beyond the most notable brain-like domain, gut endoderm derivatives and somites are also formed. We next investigated whether these tissues acquire A-P patterning as well (Figure SF7A-B). Somites in HAPs express the ubiquitous markers Foxc1 and Meox1 as well as the posterior marker Uncx and anterior marker Tbx18, suggesting A-P patterning. Similar to neural tissues, ventral (sclerotome) markers are only weakly expressed. Analysis of gut markers revealed expression of midgut/hindgut markers, showing partial A-P patterning. Together, these results show the capacity of ESC aggregates to build a unified structure with diversified tissues. Having demonstrated this capacity, in the remainder of the study we focus on mechanisms of cell fate diversification in this model.

### Hypoxia boosts the neural patterning of HAP-gastruloids along the A-P axis

The robust representation and patterning of neural cells in HAP-gastruloids is achieved by exposure to hypoxia in early commitment stages prior to elongation. We next tested whether this exposure is indeed essential to gain anterior neural fates. For this, we generated A-P gastruloids under normoxia (referred to as NAP-gastruloids, Figure 3A). NAP-gastruloids are built by assembling 2 aggregates instead of 3, since aggregates A and B are effectively the same in the absence of hypoxia. Compared to HAP-gastruloids, fewer NAP-gastruloids contained an OTX2+ region (Figure 3B-C), with the number of OTX2+ cells significantly reduced in the anterior domain (Figure 3D). PAX6 expression was more severely reduced, with less than 10% of NAP-gastruloids harboring OTX2+/PAX6+ (forebrain) cells in comparison to 80% HAP-gastruloids (Figure 3B-D). This is not due to an inherent barrier of the normoxic cells against specifying anterior neural fates, because the ‘brain’ aggregate (anterior aggregate) when cultured alone expressed and patterned OTX2+/PAX6+ cells (Figure 3E).

**Figure 3:**
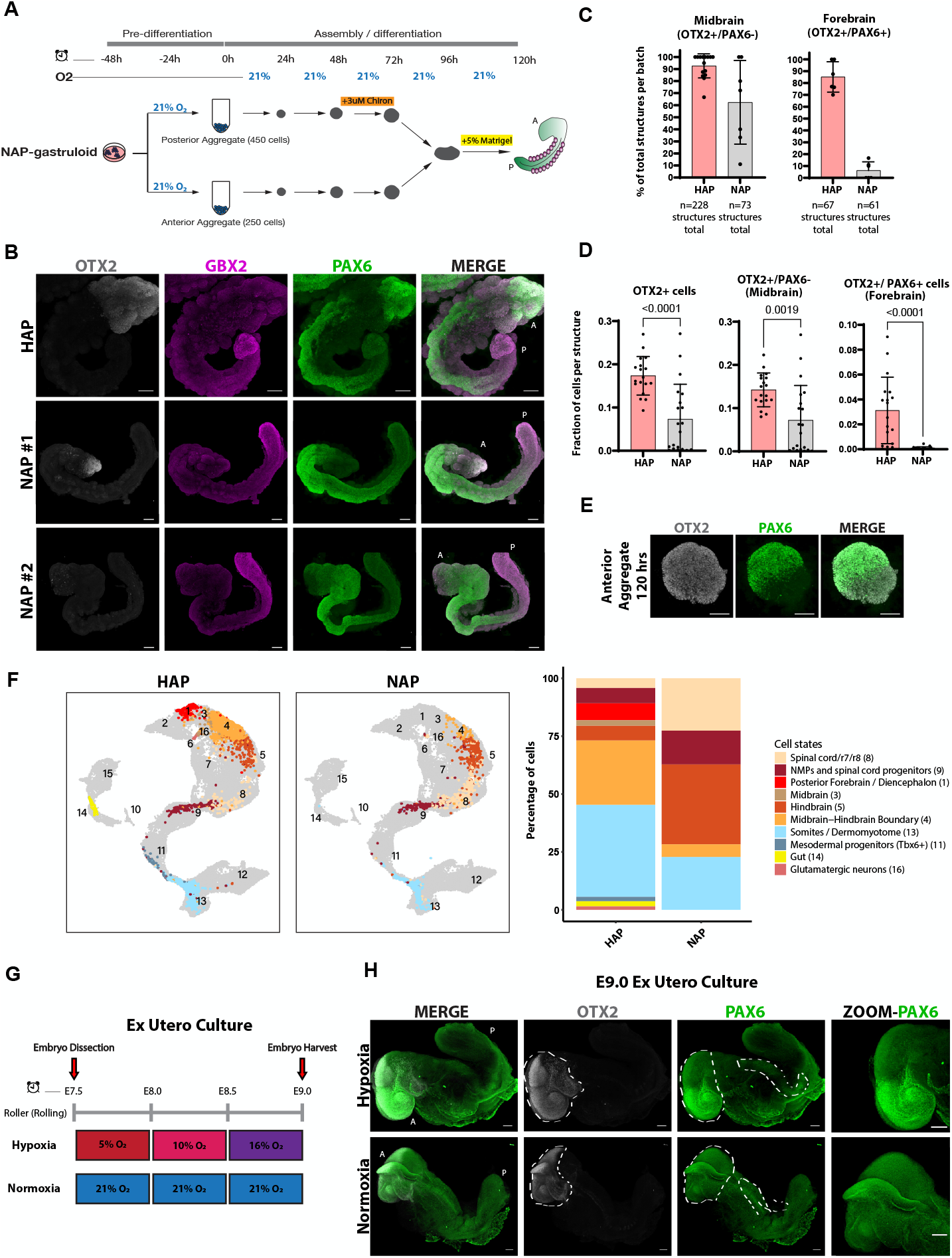
Hypoxia boosts anterior neural fates. A. Schematic illustration of the experimental pipeline for NAP-gastruloid generation. Note that only two aggregates are used, because posterior aggregates are identical without hypoxia exposure. B. IF staining of HAP-gastruloids and NAP-gastruloids for fore-/midbrain marker OTX2, hindbrain/NMP marker GBX2 and forebrain/spinal cord marker PAX6. A: Anterior, P: Posterior. Scale bars, 100 μm. C. Percentage of total structures per batch showing midbrain and forebrain formation in HAP- and NAP-gastruloids. Column heights show mean, and error bars indicate standard deviation. Each dot represents a different batch of structures. D. Fraction of cells per structure expressing the shown markers in HAP- and NAP-gastruloids. Each dot represents an individual structure. Statistical test is unpaired two-tailed t test. E. IF staining of the normoxic anterior (Chironless) aggregate at 120 hrs for fore-/midbrain marker OTX2 and forebrain/spinal cord marker PAX6. Scale bars, 100 μm. F. Left, UMAP plot of HAP- and NAP-gastruloids projected onto the embryo atlas47. Right, stacked bar plots showing cell type compositions. G. Schematic illustration of ex utero mouse embryo culture conditions. H. IF staining of E9.0 (chronological time) ex utero cultured embryos for fore-/midbrain marker OTX2 and forebrain/spinal cord marker PAX6. Dashed lines show areas of interest. A: Anterior, P: Posterior. Scale bars, 100 μm.

ScRNA-seq analysis confirmed these findings and showed that NAP-gastruloids formed no forebrain or midbrain, had a less expanded midbrain-hindbrain boundary and instead showed hindbrain and spinal cord overrepresentation (Figure 3F). Additionally, NAP-gastruloids did not form gut endoderm, as neither scRNA-seq analysis nor FOXA2 stainings showed gut derivatives (Figure 3F, S8A). We concluded that hypoxia significantly promotes anterior neural fates during cellular self-organization and differentiation, eventually boosting A-P axis neural patterning in HAP-gastruloids.

In order to test whether the developing mouse embryo uses its hypoxic microenvironment to mold neural cell fates as observed in the HAP-gastruloid model, we harvested mouse mid-gastrulation embryos and cultured these *ex utero*^32^ either in gradual decreasing hypoxia or continuous normoxia starting from E7.5 (Figure 3G). At E9.0-equivalent stage, embryos were collected and stained for PAX6 and OTX2 (Figure 3H, S8B). Both normoxic and hypoxic embryos continued to develop in culture, with hypoxic ones showing nearly 30% higher efficiency and stage-appropriate morphology by E9.0 as reported before^32^. When deeply examining the embryos we observed that normoxic culture particularly impaired forebrain formation, as normoxic *ex utero* embryos showed severely reduced PAX6 expression compared to hypoxic ones (Figure 3H). Normoxic embryos additionally showed a more restricted OTX2 expression similar to NAP-gastruloids along with spinal cord defects (Figure 3H, S8B). In contrast, hypoxic embryos showed PAX6 and OTX2 expression and neural patterning similar to HAP-gastruloids (Figure 3H, S8B). Together, these results point to hypoxia as a critical microenvironmental effector promoting proper anterior neural development and patterning in both embryo models and embryos.

### *Hif1a* KO HAP-gastruloids show spinal cord and brain defects

The cellular response to hypoxia is mainly mediated by the transcription factor HIF1a^49–51^. To genetically perturb the hypoxia response, we generated *Hif1a* knock-out (KO) ESCs by deleting the first six exons of the gene homozygously (Figure S9A-D). Using these, we then generated *Hif1a* KO HAP-gastruloids and characterized them via immunostaining and scRNA-seq (Figure 4A-F, S9E). *Hif1a* KO HAP-gastruloids showed severe neural tube (spinal cord) defects, with less than 30% showing a continuous neural tube (Figure 4A-C, S9E). This neural tube defect was not observed in *Hif1a* KO NAP-gastruloids (normoxic culture, Figure S9F), suggesting that HIF1a’s role was rescued by oxygen. Notably, *Hif1a* KO severely compromised anterior neural fates, with less than 10% forming a PAX6+ forebrain domain in comparison to 85% in the wild-type (Figure 4B-C, S9E). Similar to normoxic ex utero embryos and NAP-gastruloids, *Hif1a* KO HAP-gastruloids showed a restricted capacity to express OTX2+ midbrain cells (Figure 4B, D). These results, supported by scRNA-seq experiments (Figure 4E), suggest that hypoxia promotes anterior neural fates at least in part through HIF1a. These results are reminiscent of the phenotype of *Hif1a* KO mouse embryos which show neural tube defects and a loss of forebrain cells during their development in the hypoxic in vivo environment^33,52^. We note that, in HAP-gastruloids, anterior neural cells - in contrast to posterior neural tube/spinal cord cells - are only robustly represented in hypoxic wild-type conditions, thus underlining a necessity for HIF1a expression in anterior tissues.

**Figure 4:**
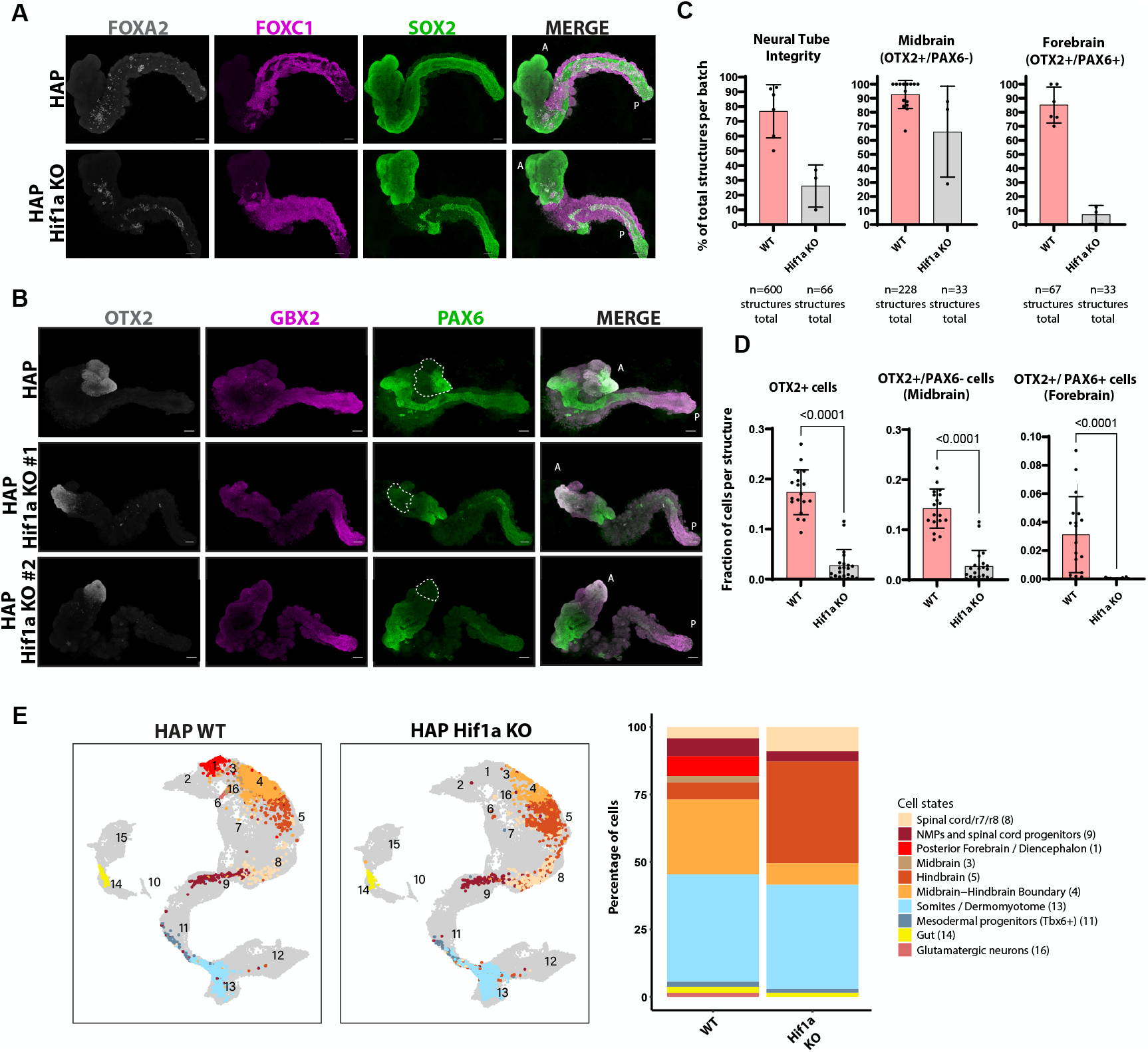
HIF1a KO compromises spinal cord and forebrain formation. A. IF staining of HAP-gastruloids and HAP-gastruloids Hif1a KO for neural marker SOX2, somitic marker FOXC1 and endoderm marker FOXA2. A: Anterior, P: Posterior. Scale bars, 100 μm. B. IF staining of HAP-gastruloids and HAP-gastruloids Hif1a KO for fore-/midbrain marker OTX2, hindbrain/NMP marker GBX2 and forebrain/spinal cord marker PAX6. Dash lines show OTX2+ regions. A: Anterior, P: Posterior. Scale bars, 100 μm. C. Percentage of total structures per batch showing midbrain and forebrain formation in wild-type and Hif1a KO HAP-gastruloids. Column height and error bars indicate mean and standard deviation. Each dot represents a different batch of structures. D. Fraction of cells per structure expressing the shown markers in wild-type and Hif1a KO HAP-gastruloids. Each dot represents an individual structure. Statistical test is unpaired two-tailed t test. F. Left, UMAP plot of wild-type and Hif1a KO HAP-gastruloids projected onto the embryo atlas47. Right, stacked bar plots showing cell type compositions.

### Perturbation of the TGFβ/BMP pathway hinders neural fate commitment and patterning in HAP-gastruloids

We next set out to investigate whether the hypoxia response in HAP-gastruloids is channeled through any of the primary pathways that are involved in brain specification during mouse embryo development, such as TGFβ and BMP, to shape neural patterning. Precise spatial and temporal regulation of the TGFβ superfamily, i.e. TGFβ, BMP and Nodal signaling, is critical for proper neural induction and subsequent brain development during gastrulation^53^. Specifically, TGFβ, BMPs and Nodal signaling must be antagonized at the anterior end of the epiblast to allow commitment towards anterior neural fates and establish the rostrocaudal boundaries in the developing brain^2,54-57^. At the same time, however, BMP activation is essential for dorsalization of the neural tube^58–60^. Rationalizing that hypoxia may modulate these pathways, we performed a series of perturbation experiments to increase or decrease their activities in HAP-gastruloids. For this, we treated the HAP-gastruloids ‘brain’ aggregate (aggregate C) with one of the following for 24h between 48-72h: TFGβ protein (2 ng/ml), TGFβ/Nodal inhibitor SB-431542 (10 uM), BMP inhibitor Noggin (100 ng/ml) (Figure 5A), BMP4 (10 ng/ml) or BMP7 (25 ng/ml) (Figure S10A). The applied concentrations are commonly used in liver and brain organoid generation^45,61-64^.

**Figure 5:**
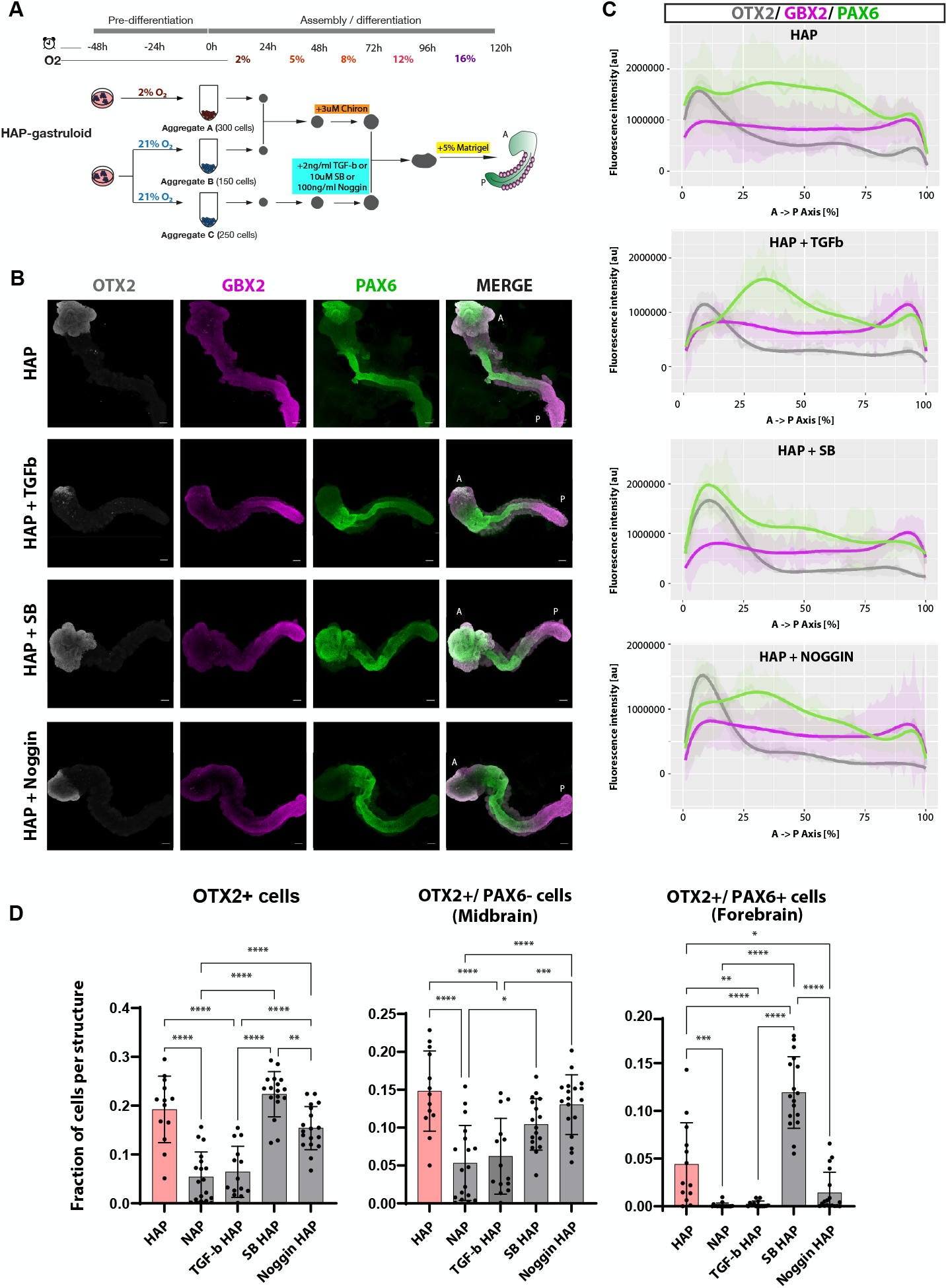
Brain formation in HAPs is hindered by TGF-b overexpression and BMP inhibition. A. Schematic illustration of the experimental pipeline for treating HAP-gastruloids with TGFβ, SB or Noggin. B. Left, IF staining of HAP, HAP+TGFβ, HAP+SB and HAP+Noggin for fore-/midbrain marker OTX2, hindbrain/NMP marker GBX2 and forebrain/spinal cord marker PAX6. A: Anterior, P: Posterior. Scale bars, 100 μm. Right, Localization of OTX2, GBX2 and PAX6 signals along the A-P axis of each condition. Lines show mean values from 10 individual structures. C. Localization of OTX2, FOXG1 and PAX6 signals along the A-P axis of HAP, HAP+TGFβ, HAP+SB and HAP-+Noggin. Lines show mean values from 10 individual structures D. Fraction of cells per structure expressing OTX2+, OTX2+/PAX6-(midbrain) and OTX2+/PAX6+ (forebrain) cells in HAP, HAP+TGFβ, HAP+SB and E. HAP+Noggin. Column height and error bars indicate mean and standard deviation. Each dot represents an individual structure. HAP-gastruloids: a total of 13 individual structures from 1 biological replicate were used. HAP+TGFβ: a total of 14 individual structures from 2 biological replicates were used (7 structures/biological replicate). HAP+SB: a total of 17 individual structures from 2 biological replicates were used (8-9 structures/biological replicate). HAP+Noggin: a total of 18 individual structures from 2 biological replicates were used (9 structures/biological replicate). Statistical test is one-way ANOVA with multiple testing correction. P-values **** p<0.001, ** p<0.005, *p<0.05

Fluorescence staining and quantifications showed that TGFβ supplementation significantly reduced midbrain and forebrain formation (fewer OTX2+/PAX6+ cells, Figure 5B-D, S11), whereas TGFβ inhibition overcommitted cells towards forebrain and compromised regionalization (more OTX2+/PAX6+ cells, Figure 5B-D, S11). BMP4 supplementation completely altered HAP-gastruloid morphology and remained inconclusive (Figure S10B). Notably, BMP inhibition and BMP7 supplementation severely reduced forebrain but not midbrain commitment (Figure 5B-D, S10C-D), indicating that TGFβ and BMP pathways distinctly regulate mid- and forebrain regionalization. These results suggest that hypoxia may enable the commitment to diverse anterior neural fates at least in part by modulating TGFβ and BMP expression levels.

### Anterior forebrain development in HAP-gastruloids with WNT inhibition

Despite robust commitment to anterior neural cells in HAP-gastruloids, the absence of the most rostral forebrain markers remained a major limitation to fulfilling the full complexity of the brain (Figure 2G, 6A). During organogenesis, around E11, the anterior forebrain forms the telencephalon, which gives rise to the cortex, the center of mammalian cognition. Although telencephalon expands around E10, the telencephalic primordium is specified at 8.5, marked by the expression of the forkhead transcription factor gene FOXG1 ^65,66^ (Figure 6B). Given that during gastrulation the most anterior end of the epiblast (future rostral forebrain) is exposed to signals that inhibit WNTs emanating from the extraembryonic tissues ^67–69^, we rationalized that residual WNT activity or insufficient WNT inhibition in HAP-gastruloids could hinder anterior forebrain formation. We thus tested whether exogenous WNT inhibition (WNTi) could further promote brain anteriorization by treating the ‘brain’ aggregate (aggregate C) with the WNT inhibitor XAV-939 (Figure 6C, HAP-gastruloids+XAV’ condition, treatment window is 48-72h). Remarkably, WNTi under hypoxia induced robust, but not ectopic, FOXG1 expression and the majority of HAP-gastruloids+XAV structures expressed FOXG1 in a similar pattern to the E8.75 embryo (Figure 6D-F, S12A). The total fraction of OTX2+ cells remained similar between HAP-gastruloids and HAP-gastruloids+XAV, however the spatial organization of the brain compartments was more advanced in HAP-gastruloids+XAV, with midbrain-, posterior forebrain-, and anterior forebrain-like regions clearly distinguishable from each other (Figure 6E). ScRNA-seq validated these results, showing the expression of the anterior forebrain markers Lhx2, Six3, Fezf1 and Foxg1 in HAP + XAV condition (Figure 6G, S12B). Surprisingly, we also detected Nkx6.1 and Olig2 expression, suggesting ventralization of the spinal cord (S12B).

**Figure 6:**
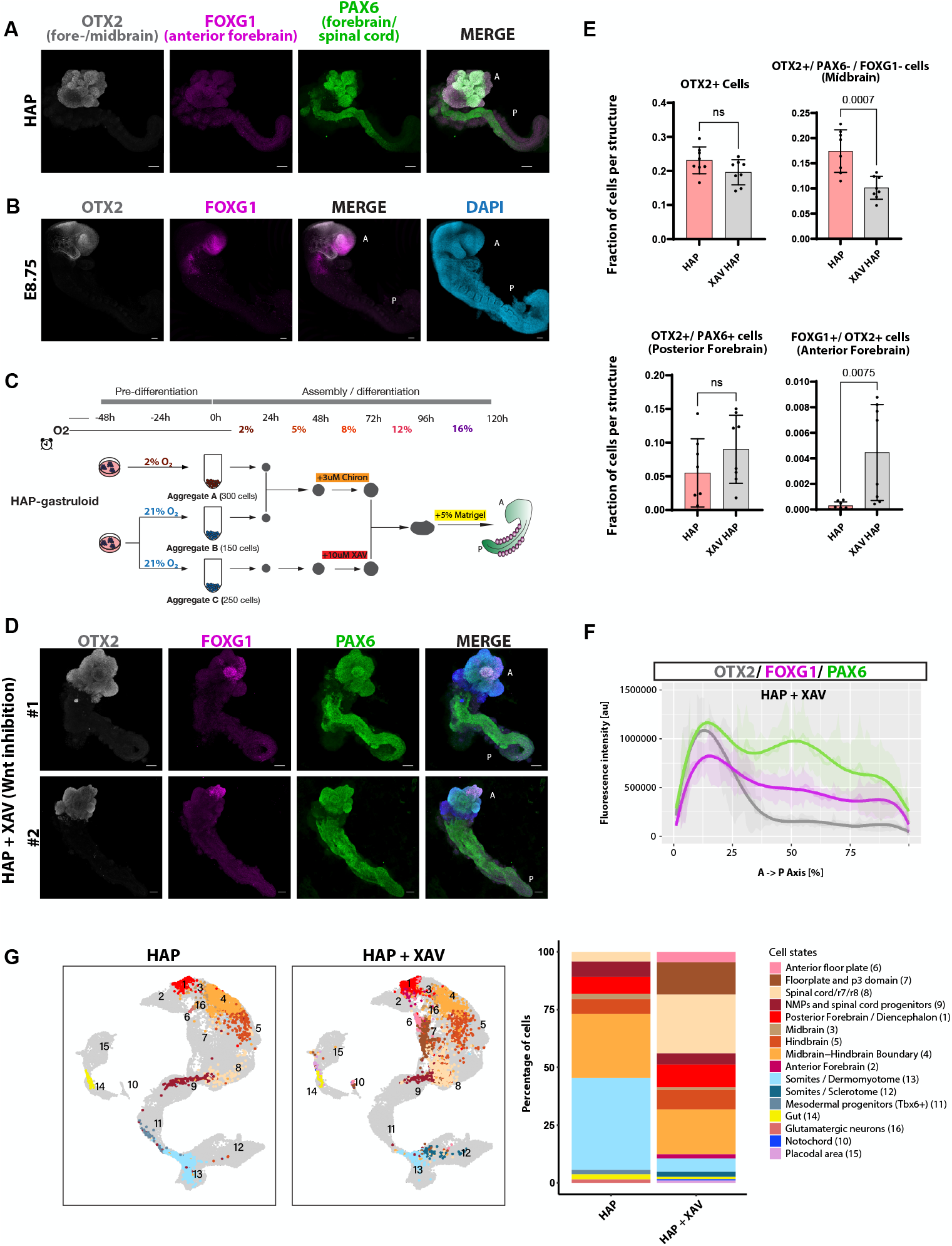
Wnt inhibition induces anterior forebrain. A. IF staining of HAP-gastruloids for the fore-/midbrain marker OTX2, anterior forebrain marker FOXG1 and forebrain/spinal cord marker PAX6. A: Anterior, P: Posterior. Scale bars, 100 μm B. IF staining of E8.75 mouse embryo for the shown markers. Scale bars, 100 μm C. Schematic illustration of the experimental pipeline for treating HAP-gastruloids with XAV. D. IF staining of HAP-gastruloids + XAV for fore-/midbrain marker OTX2, anterior forebrain marker FOXG1 and forebrain/spinal cord marker PAX6. A: Anterior, P: Posterior. Scale bars, 100 μm E. Fraction of cells per structure expressing the shown markers in HAP-gastruloids with or without XAV treatment. Column height and error bars indicate mean and standard deviation. Each dot represents an individual structure. Statistical test is unpaired two-tailed t test. F. Localization of OTX2, FOXG1 and PAX6 signals along the A-P axis of HAP-gastruloids+XAV. Lines show mean values from 8 different structures. G. Left, UMAP plot of HAP- and HAP+XAV projected onto the embryo atlas47. Right, stacked bar plots showing cell type compositions.

Finally, in order to test whether TGFβ or WNT inhibition can substitute for hypoxia in the formation of a well-developed and patterned brain domain in HAP-gastruloids, we treated NAP-gastruloids (normoxic gastruloids, Figure 3A) with either SB or XAV or both (Figure S13A). In line with this, TGFβ inhibition in NAP-gastruloids yielded an expanded OTX2+ midbrain domain, but not OTX2+/PAX6+ forebrain (S13B), while WNT inhibition of NAP-gastruloids yielded an OTX2+ midbrain domain, occasionally (40%) OTX2+/PAX6+ posterior forebrain, but no FOXG1+/OTX2+ anterior forebrain (Figure S13C). Inhibition of both TGFβ and WNT pathways yielded structures with OTX2+/PAX6+ domains, but none with FOXG1+/OTX2+ anterior forebrain (Figure S13D). Based on these results, we concluded that external cues do not substitute hypoxia treatment and that hypoxia-driven cell-intrinsic regulation of TGFβ in combination with exogenous WNT inhibition are necessary for advanced anterior neural patterning in HAP-gastruloids.

## DISCUSSION

The developing embryo does not emerge from the simple assembly of distinct parts. It is only through the study of stem cell-derived aggregates that the concept of modularity begins to emerge. While the formation of HAP-gastruloids does not serve as a direct replica of the embryonic process, it provides valuable insights into the principles of cellular interactions within multicellular systems. Importantly, we emphasize that HAP-gastruloids were not developed to serve as a perfect mimic of the embryo. Instead, our investigation of hypoxia as a critical microenvironmental factor led to the development of HAP-gastruloids, which in turn facilitated the study of interactions between different cellular populations within these aggregates.

Similar to other in vitro systems, such as ESC cultures and gastruloids, HAP-gastruloids selectively recapitulates key aspects of embryogenesis. HAP-gastruloids open new avenues for investigating cellular processes that are otherwise inaccessible in embryos. In our model, we chose to use ESCs without the support of extraembryonic cells, simplifying the system and enabling us to specifically focus on epiblast-driven decision-making. The assembly approach, when combined with hypoxia, recreates the natural morphogen gradients in the epiblast, ultimately yielding a unified embryo model with anterior-posterior (A-P) patterning characteristic of early development.

Often referred to as a detrimental element in somatic tissues, hypoxia is an integral part of mammalian development. In addition to inducing a metabolic shift away from oxidative phosphorylation and towards glycolysis, hypoxia can affect the transcriptional profile through HIF1a and/or by altering the activity of histone and DNA demethylases^31,70^. The hypoxia-driven cell type diversification and compartmentalization in our system is largely mediated by HIF1a, as *Hif1a* KO HAP-gastruloids show compromised neural commitment. Our findings in the HAP-gastruloids model are consistent with the embryonic phenotypes of *Hif1a* and *Hif2a* KO mice that show neural tube and brain defects, including perturbation of forebrain patterning^33,52^.

Additionally, the HAP-gastruloids model now allows spatial dissection of such phenotypes, by e.g. creating anterior-only KOs.

We show that the benefit of hypoxia is negated upon perturbation of TGFβ levels, suggesting that TGFβ expression is oxygen-sensitive in our system similar to others^71,72^. It is plausible that TGFβ is directly regulated by HIF1a or through mechanisms independent of HIF1a or through other effectors downstream of HIF1a, eventually resulting in TGFβ inhibition. Additionally, considering BMP signaling is necessary for dorsal patterning, it is plausible that hypoxia regulates selective BMPs for dorsal patterning of the forebrain, as BMP inhibition disrupts dorsal forebrain (PAX6 expression) but no midbrain formation. However, whether BMPs are regulated by hypoxia and if this regulation is driven by HIF1a remains to be seen in further studies. In summary, our findings build on and uniquely expand decades of key research on the effects of hypoxia by revealing its key influence in the developing embryo and our embryo model.

Beyond investigating hypoxia, HAP-gastruloids offer a unique model to investigate self-organizing properties and patterning events beyond the findings presented here. We show that the assembled aggregates that make up the HAP-gastruloids form a homochronic entity at 120h. We also show that the ‘brain’ aggregate distinctly forms the anterior neural ‘brain-like’ region of HAP-gastruloids, while the other Chiron-treated aggregates form the posterior tissues. Notably, it is the integration of these anterior and posterior regions and the interaction between them that forms a continuous neural tube and an anterior brain-like domain organized regionally along the A-P axis. Overall, HAP-gastruloids is a unique embryo model that can be used to address key open questions otherwise concealed by the uterine microenvironment.

## ACKNOWLEDGMENTS

We thank members of the Bulut-Karslıoğlu Lab and Matthew Kraushar, Sigmar Stricker, and Ludovic Vallier for critical feedback; MPIMG facilities, especially microscopy, for excellent service and discussions. We thank Can Aztekin, Kelly Hu, and Marion Leleu for help with scRNA-seq. We thank Maria Walther, Susanne Lade, Astrid Grimme, Cordula Mancini, Birgit Romberg, Shelby Baumgartner, Edda Einfeldt for assistance. Work in the Hanna Lab is supported by MBZUAI-WIS Program grant, a Minerva Stiftung grant, and ERC-COG-2022 #101089297– ExUteroEmbryogenesis. Work in the Kretzmer Lab is supported by the Hasso Plattner Institute and the Max Planck Society. Work in the Bulut-Karslioğlu Lab is supported by the Max Planck Society, and Humboldt Foundation (Sofja Kovalevskaja Award), the European Research Council (ERC) under the European Union’s Horizon 2020 research and innovation programme (ERC grant agreement no. 101117421, ‘DOR CODE’’).

## AUTHOR CONTRIBUTIONS

Conceptualization: AB and AB-K. Methodology: AB and AB-K. Investigation: AB, HÖÖ, IB. Formal analysis: IK, PAO, RB. Visualization: AB, IK, RB. Writing - original draft: AB and AB-K. Supervision: JHH, HK, AB-K. Funding acquisition: JHH, HK, AB-K.

## METHODS

### Animal experimentation

All animal experiments were performed according to local animal welfare laws and approved by local authorities (Landesamt für Gesundheit und Soziales). Mice were housed in individually ventilated cages and fed ad libitum. For mouse dissection the uterus was initially isolated at the desired stage, was cut between the decidua and placed with a strainer spoon (Catalog # TL85.1, ROTH) into a dish containing embryo medium (DMEM/F-12 without phenol red (Thermo Fisher Scientific, cat# 21041025) + 10% FBS). Muscle tissue around the decidua was removed with forceps (Dumont #55 Forceps, catalog #11255-20, FST) under the stereomicroscope. Decidua was opened and embryos were carefully removed with forceps. The remaining decidua tissue was cleaned and the amnion was carefully opened. Embryos were placed in a dish containing embryo medium with a pipette and kept on ice for fixation.

### Mouse ESC culture

F1G4 and FK223 (received from Bernhard Herrmann Lab, FK233 is the reporter cell line) and KH2 (received from Alexander Meissner Lab) mouse ESC lines were used. Cells routinely tested negative for mycoplasma, were however not authenticated. ESCs were routinely maintained on 6 cm plates (Corning, 430166) gelatinized with 0.1% gelatin (1:20 dilution of 2% gelatin (Sigma, G1393) in tissue-culture grade H_2_O) and coated with mitotically inactive primary mouse embryo fibroblasts (3-4×10^4^ cells/cm^2^). Cells were cultured in Knockout Dulbecco’s Modified Eagle’s Medium (KO-DMEM) (Thermo Fisher Scientific 10829-018) supplemented with 15% FBS (Thermo Fisher Scientific, Cat# 10438026), 1% GlutaMAX (GIBCO, 35050038), 1% nucleosides (EmbryoMax Nucleosides (100x, Merck, ES-008-D), 0.2% beta-mercaptoethanol (Thermo Fisher Scientific, 21985023), Penicillin-Streptomycin (10,000 U/mL) (GIBCO, 15140122), and 1000 U/mL homemade LIF and grown at 37°C in a 21% O_2_ and 5% CO_2_ incubator unless otherwise specified. Hypoxic ESCs were grown in 2% O_2_. mESCs were split every second day with a dilution suitable to the proliferation velocity (between 1:5 and 1:9). mESC+LIF medium was refreshed daily. For splitting, media was aspirated and cells were washed once with PBS and dissociated with TrypLE (Thermo Fisher Scientific 12604-021) for 5-10 min at 37°C. TrypLE was neutralized by 3 ml mESC+LIF and cells centrifuged for 5 min at 1000 rpm, after which the pellet was resuspended in mESC+LIF. For freezing of mESCs, cell pellets were resuspended in mESC medium with 20% FCS, and mixed in a 1:1 ratio with mESC freezing medium. Cells were frozen down overnight in the –80 °C and transferred to liquid nitrogen the next day.

### Optimization of O_2_ implementation in TLS model

We first systematically tested different oxygen concentrations before and during TLS formation (Figure S1, conditions I-IX). Early and continuous exposure to 2% O_2_ (conditions II-III) boosted gut endoderm formation but compromised the axial elongation and overall area, which were rescued when the structures were exposed to normoxia in the last 48h (condition V). These results suggested that a combination of low and high oxygen levels might be needed for optimal cell type composition and morphology. We next integrated early low and late high oxygen levels, with a gradual increase in between during TLS generation (conditions VI-IX). These new setups led to the formation of well-elongated structures, which, however, exhibited short neural tubes and disorganized anterior regions. Concurrently, exposing cells in low oxygen before differentiation (condition IX) promoted gut endoderm formation, but mildly compromised morphology. In comparison, pre-exposing cells in normoxia before differentiation did not compromise morphology, but also did not form gut endoderm (condition VIII). We concluded that a single-aggregate system is incompatible with consistent morphological outcomes and enhanced cell type composition.

### HAP-/NAP-gastruloid generation

mESCs were cultured in either hypoxia and normoxia. At the onset of HAP-gastruloid generation, normoxic and hypoxic mESCs were trypsinized, washed with mESC medium and resuspended as single cells. After MEF depletion, mESCs were pelleted and resuspended in PBS. After another wash, cells were resuspended in pre-warmed NDiff227 (Takara, XA0530) and counted with Countess 3 (Invitrogen, 16842556). Cells were plated in a 96-U-bottom, ultra-low attachment plate (Costar, 7007) to from three distinct aggregates: aggregate A (300 cells), aggregate B (150 cells) and aggregate C (250 cells).

Aggregates A and B were fused at 24h and treated with 3 uM Chiron at 48h. Aggregate C was grown separately without being treated with Chiron and at 72h fused with the other aggregates. At 96 h, 5% matrigel (corning/BD, 356231) was added. At 120 h, structures were collected for further processing. Structures were grown in a hypoxic incubator at 37°C and 5% CO2 in a gradually increasing oxygen setup. NAP-gastruloids were generated following the same procedure but cultured in 21% O2.

### Generation of *Hif1a* KO mouse ES cells

*Hif1a* KO gene was knocked out with CRISPR/Cas9 gene editing tool. More specifically the first six exons of the gene were removed by gRNAs targeting upstream of exon 1 (TCAGAGAACTCTATAGAGAGCCG) and downstream of exon 6 (TTGTACGTCCACTGTATCCAAAG). Grna sequences were cloned into the pX330A-1x2 plasmid. Wild-type F1G4 cells were nucleofected with the plasmids using the Lonza 4D Nucleofector. After 48 h, cells were single-cell sorted into 96-well plates using the BD FACSAria Fusion (Software v8.0.1). 8-10 days later clones were selected and expanded. After expansion DNA was extracted for each clone and screened using two primer pairs: Primer pair 1: Knockout Primers, amplicon size 767 bp, amplified only if KO is successful. F: 5′-GAGAGGTTTGGCAAGACGGAT-3’ R: 5′-TGTGCAAGGGATGTTGCCTAA-3’ Primer Pair 2: Control Primers, amplicon size 653 bp, amplified only if KO is unsuccessful. F: 5′-AGTGGCTAAGGAAGTAAGCACC-3’ R: 5′-ACATTGTGGGGAAGTGGCAA-3’

### DNA extraction

Cells were grown until confluent enough. Lysis buffer was added and cells were incubated overnight at 37°C. Next day ice cold NaCl/EtOH was added for 2h/RT, until proper precipitating DNA was visible. DNA was then washed 3x with 70% EtOH and air dried for 15-20 minutes. Finally, DNA was resuspended with the desired volume of H2O by overnight incubation at 37°C. For subsequent genotyping PCR usually 1 or 2 ul DNA was used as a template.

### RNA extraction and RT-qPCR

Total RNA was extracted using the QIAGEN RNeasy kit (QIAGEN, 74004), following the manufacturer’s instructions and 5 µg of RNA was used as input to generate complementary DNA (cDNA) with the High-Capacity cDNA Reverse Transcription Kit (Thermo Fisher Scientific, 4368814). cDNA was used to perform real-time quantitative PCR (qPCR) using primers designed to amplify different exons of *Hif1a* using KAPA SYBR FAST qPCR Master Mix (2X) ABI Prism (Thermo Fisher Scientific, KK4617) on the QuantStudio 7 Flex Real-Time PCR System (Applied Biosystems) thermal cycler. In RT-qPCR experiments, data represent log_2_FC over *mTbp*.

Primers used:

Exon2_3:

F: 5’-TCTCGGCGAAGCAAAGAGTC-3’

R: 5’-AGCCATCTAGGGCTTTCAGATAA-3’

Exon5_6:

F: 5’-GATGACGGCGACATGGTTTAC-3’

R: 5’-CTCACTGGGCCATTTCTGTGT-3’

Exon10_11:

F: 5’-ACCTTCATCGGAAACTCCAAAG-3’

R: 5’-CTGTTAGGCTGGGAAAAGTTAGG-3’

mTbp:

F: 5’-CCTTGTACCCTTCACCAATGAC-3’

R: 5’-ACAGCCAAGATTCACGGTAGA-3;

### Nucleofection of ESCs

Nucleofection was performed according to Amaxa UD-Nucleofector Protocol provided by the company. Amaxa 4D Nucleofector kit (Lonza V4XP-3024) and 4d-Nucleofector unit were used.

### Ex utero embryo culture

Ex utero culture was performed as described in^32^. All embryos were dissected at E7.5, immediately moved to roller culture, grown in EUCM medium and harvested 1.5 days later (chronological time E9.0). During roller culture, two different O_2_ conditions were tested as indicated in the figures.

### Inhibitor/Growth Factor treatments

For exogenous WNT activation: 3 uM Chiron (CHIR99021, Sigma, SML1046), for exogenous TGFβ supplementation: 2 ng/ml TGFβ (Biotechne, 240-B-010), for TGFβ and Nodal inhibition: 10 uM SB 431542 (Tocris, 1614/10), for BMP signalling inhibition 100 ng/ml Noggin (Recombinant Human Noggin/Fc Chimera, CF 50 ug, R&D Systems, 3344-NG-050), for exogenous BMP4 supplementation 10 ng/ml BMP4 (R&D Systems, 314-BP-500/CF), for exogenous BMP7 supplementation 25 ng/ml BMP7 (Thermo Fisher, PHC9544), for WNT inhibition: 10 uM XAV939 (MedChemExpress, HY-15147).

### Immunostaining

Whole mount immunofluorescence was performed as described in^10^. Primary antibodies used in this study are: OTX2 (Polyclonal Goat IgG, R&D Systems, AF1979), PAX6 (mouse hybridoma, DSHB (concentrated)), PAX6 (Polyclonal Rabbit IgG, BioLegend, 901301), GBX2 (Polyclonal Rabbit IgG, Proteintech, 21639-1-AP), FOXG1 (Monoclonal Rabbit IgG, Abcam, [EPR18987] ab196868), FOXC1 (Monoclonal Rabbit IgG, Abcam, [EPR20685] ab227977), FOXA2 (Polyclonal Goat IgG, R&D Systems, AF2400), SOX2 (Monoclonal mouse IgG2a, Santa Cruz, sc-365964). All primary antibodies were used in a dilution 1:200, apart from GBX2 (1:100) AND PAX6 (mouse hybridoma, 1:50). Secondary antibodies used in this study include: Donkey anti-Goat IgG (H+L) Cross-Adsorbed Secondary Antibody, Alexa Fluor™ 647 (Thermo, A21447), Donkey anti-Mouse IgG (H+L) Highly Cross-Adsorbed Secondary Antibody, Alexa Fluor Plus 488 (Thermo, A32766) and Donkey anti-Rabbit IgG (H+L) Highly Cross-Adsorbed Secondary Antibody, Alexa Fluor 546 (Thermo, A10040). All secondary antibodies were used in a dilution 1:250.

### Hybridization Chain Reaction Fluorescence In Situ Hybridization for RNA (HCR RNA FISH)

HCR RNA–FISH was performed according to the protocol from Molecular Instruments.

### Tissue clearing

Prior to imaging, mouse embryos and AP-gastruloids were cleared with RIMS (Refractive Index Matching Solution). To this end, samples were washed twice with PBS for 10 min, post-fixed in 4% PFA for 20 min and washed three times with 0.1M phosphate buffer (PB, 0.025 M NaH_2_PO_4_ and 0.075 M Na_2_HPO_4_, pH 7.4). Clearing was performed by incubation in RIMS (133% w/v Histodenz (Sigma, D2158 in 0.02M PB) on a rocking platform at 4°C for at least one day. Occasionally, HAP-gastruloids were embedded in agarose before cleared with RIMS. To this end, 1.5% low melting point (LMP), analytical grade (Promega, V2111) agarose was prepared in PBS, incubated at 80°C for 15 min, and cooled down to 37°C n in a thermomixer. The pipette was set to 20 µl and HAP-gastruloids were stabilized on the ibidi plate with a drop of LMP agarose for 5 min until the agarose was dry.

### Image acquisition

Confocal images were acquired on a Zeiss LSM 880 confocal running under ZEN Black 2.3 (Zeiss Germany). All structures were captured in z-Stacks with an Airydetector in FastAiryMode. The individual structures were recorded in tile scans using a Plan-Apochromat 20x and a numerical aperture of 0.8. The lateral (xy) resolution was 0.25 µm per pixel, furthermore a distance of 10 µm was chosen for the axial (z) resolution in order to image an average nucleus diameter and to avoid oversampling of signals for later expression analyses. The airy data were processed and prelayd for several downstream analyses.

Wide field images were acquired using the Zeiss Celldiscoverer 7 running under Zen Blue 3.7 (Zeiss Germany). Structures were captured in z-stack with an Axiocam 712. An 5x PlanApochromat with a numerical aperture of 0.35 and a 3×3 camera binning achieved a lateral resolution of 2.09 µm per pixel, also here the slice height was captured with 10 µm. Each captured structure was located in a single mold of 96-well ultra-low attachment plate and served as a single field of view.

### Image Processing

Maximum Intensity Projections were used to quantify the expression of key markers along the morphological axes. To generate these in batches from raw 3-dimensional confocal data, we used a self-made script in ZEN 3.8 blue in the module “Open Application Development” as well as functions of the module “Advanced Processing”. Batched airyscan processing of confocal data was performed on a dedicated workstation running under ZEN Black v2.3 (Zeiss, Germany).

### Image Analysis

#### Single cell measurements of fluorescence intensity and % positive cells

Analysis was performed using Zen Blue module “Image Analysis” running under v3.5 (Zeiss, Germany). All confocal data was processed with the identical fully automated pipeline using a classical segmentation algorithm. Herein, first the nuclear counterstain was smoothed, then the background was subtracted. Objects were identified by applying an intensity threshold to the foreground, followed by classical water shedding and a size filter between 100-1000 µm^2^. On average we recovered 5-10×10^5^ single nuclei per quantified structure. Fluorescence intensities and geometries of the identified objects were quantified in ZEN, Positivity threshold for individual markers were identified by first visualizing the distribution of mean intensity data in histograms, then selecting the inflection point distinguishing negative and positive populations in the bimodal distributions. The data reported in the figures show either % cells per structure, or % structures per batch.

#### Morphometrical and A-P analyses

Custom ImageJ/Fiji macros were used. In summary, primary objects were determined on counter stainings using variance-based thresholds. The primary object was used in the pipeline to determine an anterior-posterior (AP) axis using a custom-built algorithm via the ‘skeletonize’ function. The AP axes were binned to 50 intervals. Furthermore, the axis was used to ‘straighten’ the entire structure for quantifications along the AP axis. All axes were adjusted to a length of roughly 1000 pixels to enable a direct comparison over multiple conditions. For easier visualization we fitted a 6th order polynome (bold lines) over the averaged results (opaque) and standard deviation (more opaque).

### Single-cell RNA-seq

#### AP-gastruloids

Single-cell RNA-sequencing (scRNA-seq) experiments were performed using Parse Biosciences whole transcriptome kit. Samples were fixed according to user manual (Evercode Cell Fixation kits Reagents CF100) and enhancer (CF200)). Briefly, approximately 20 structures per condition were collected in a 1.5 ml protein low binding tube filled with cold PBS. After two subsequent washes with cold PBS, 300 ul TrypLE was added in each tube and the suspension was incubated for 20 minutes at 37°C, with pipetting up and down every 10 minutes till all cells were dissociated. The cell suspension was resuspended in 600 ul PBS, spinned down (4°C, 500g, 5min) and the pellet was resuspended in 1 ml PBS. After counting the desired number of cells (between 10^5-10^6cells) each sample was transferred in a new Protein Low Binding 1.5 ml tube and centrifuged at 4°C, 500xg, 5 min. Pellet was resuspended in 187.5 ul Prefixation Master Mix (see user manual for every Master Mix preparation), strained into a new 1.5 ml tube and mixed with 62.5 μL of Cell Fixative Master Mix. After a 10-minute incubation on ice, 20 μl of Permeabilization Solution were added and the sample was incubated on ice for 3 minutes. After incubation sample was mixed with 250 ul Fix and Perm Stop Buffer and spinned down (4°C, 500xg, 5 min). The pellet was resuspended in 70 ul Cell Storage Master Mix and strained into a new 1.5 ml tube. Fixed cells were counted and stored at −80°C. Library preparation was performed according to instructions (Parse Biosciences, Evercode Whole Transcriptome Version 3 kit, WT100/WT200). Sequencing was performed on Novaseq.

TLS dataset is a reference dataset generated in our lab following manufacturer’s recommendations for Chromium single cell kit (10x Genomics) prior to HAP-/NAP-gastruloid samples.

### Bioinformatics Methods

Statistical analyses, computations and plots were executed using R version 4.4.1’ “Race for Your Life”^73^. For single-cell analysis, the Seurat package version 5.1.0 was utilized^74^. Data distributions were visualized through boxplots using the ggplot2 package^75^. In these boxplots, the central line represents the median, the box defines the interquartile range (IQR) from the 25th to 75th percentile and the whiskers extend to 1.5 times the IQR. Points outside this range are identified as outliers.

### Pre-processing

scRNA-seq data were pre-processed following Parse Biosciences guidelines. Following alignment, UMI and barcode quantification took place. For each sample, 15 objects (3 runs × 5 lanes) were merged into a single Seurat object. These four sample-level objects were subsequently merged into one combined Seurat object for downstream analysis. To remove doublets, low-quality or dying cells, cells with fewer than 300 or more than 7,000 detected genes, fewer than 700 or more than 30,000 total counts, or more than 5% mitochondrial counts were removed from the analysis. After filtering, the data were log-normalized using NormalizeData(), variable features were identified with FindVariableFeatures(), and the data were scaled with ScaleData() using Seurat with default parameters.

### TOME reference

The “Trajectories Of Mouse Embryogenesis (TOME)” data were used to construct a scRNA-seq reference spanning embryonic days E7.25 to E10.5^46^. To this end, stage-specific Seurat objects were loaded and merged into a single Seurat object. Gene identifiers were mapped to gene symbols using the mapIds() function from the org.Mm.eg.db package. Genes without corresponding symbols were retained using their Ensembl IDs. To ensure unique gene representation, expression counts for duplicated symbols were summed.

Next, cells were split by sequencing batch (orig.ident), and each batch was independently normalized, scaled, and processed for principal component analysis (PCA) using a shared set of 3,000 highly variable genes selected with SelectIntegrationFeatures(nfeatures = 3000). Reciprocal PCA (RPCA) integration anchors were then computed across batches using FindIntegrationAnchors(reduction = “rpca”, dims = 1:30, k.anchor = 5, k.filter = 50), and data were integrated with IntegrateData(dims = 1:30). The integrated assay was subsequently scaled, and dimensionality reduction was performed using RunPCA(npcs = 50).

The TOME reference (E7.25-10.5) comprises 81 distinct cell state clusters. To better annotate the neural cell states, we extracted the forebrain/midbrain and hindbrain clusters and re-clustered them using Seurat. In brief, after scaling and PCA, subclustering was performed with FindNeighbors(dims=1:50) followed by FindClusters(resolution=0.01) for each cluster. The resulting subclusters were annotated based on marker gene expression and classified as forebrain, midbrain, unassigned, and hindbrain 1-3. Cell state-specific marker genes were identified using FindAllMarkers(only.pos = TRUE, min.pct = 0.05, logfc.threshold = 0.25) and filtered to retain genes with an adjusted *p*-value < 0.05.

### OMG reference

The “Ontogeny of Mouse, Graphed” (OMG) reference used in this study was constructed from scRNA-seq data spanning mouse embryonic days E3.5–13.5 ^47^. Cell-level metadata were retrieved and filtered to include only embryos from E8.0– 10.0. Count matrices and cell annotations were downloaded for the relevant runs (run_4, run_15, run_17_sub1, and run_17_sub2), together with a shared gene annotation table. For each dataset, the sparse matrices were loaded, annotated with gene and cell identifiers, and subset to include only cells present in the filtered metadata. Individual Seurat objects were created for each run using CreateSeuratObject(). Cells were quality-filtered by retaining only those with 1,500– 3,000 detected genes and 2,500–6,000 total counts. After filtering, each developmental stage was downsampled to 15,000 cells to ensure balanced representation across stages. The downsampled datasets were then merged into a single Seurat object. Gene identifiers were mapped to gene symbols using the provided annotation table, and genes with non-unique symbols were excluded to avoid redundancy. The final counts matrix was used to reconstruct the Seurat assay to maintain consistency between metadata and expression layers.

As HAP cells were predicted to correspond to E8.5 in the developmental stage analysis with TOME, the OMG reference was restricted to embryos from E8.0–E9.75. The resulting dataset was normalized, highly variable genes were identified, and scaling was performed using Seurat with default parameters. After normalization and scaling, subclustering with FindClusters(resolution = 0.1) identified three distinct facial mesenchyme cell clusters.

### Cell state annotation and projection

Cell state, developmental stage, and somitic count labels were transferred using FindTransferAnchors(dims = 1:50, reference.reduction = “pca”) followed by MapQuery(reference.reduction = “pca”). To ensure robust comparisons across datasets, quantitative filters were applied based on label prediction confidence and cell state representation. For each dataset, the 20^th^ percentile of the prediction scores was calculated for each label, and cells with scores below these thresholds were excluded to remove low-confidence annotations. In addition, within each sample, cell states represented by fewer than 25 cells were excluded.

For visualization, UMAPs were generated for the relevant TOME and OMG subsets. Each reference subset included only the cell states present in the corresponding gastruloid dataset, as determined through label transfer and filtering. UMAP embeddings were computed using RunUMAP(dims=1:30, return.model=TRUE) on scaled and PCA-transformed subsets. The gastruloid samples were then projected onto the reference UMAPs using MapQuery(reference.reduction = “pca”, reduction.model = “umap”), following anchor identification as described above. To compare the expression of marker genes, expression values were scaled within the analyzed subset of cells, restricted to the relevant samples and cell states. For each sample and cell state, the percentage of expressing cells was calculated as the proportion of cells with detectable expression relative to the total number of cells in that condition and state.

### Prediction of developmental stage and somite count

In addition to assigning each cell to the developmental stage with the highest prediction score, a continuous ‘pseudostage’ was computed as the weighted sum of all stage scores. For this calculation, the OMG stage label ‘E8.0-8.5’ was converted into 8.25, and the TOME labels ‘E8.5a’ and ‘E8.5b’ were merged into a single stage, 8.5. Somite count was predicted using the same weighted-sum approach, providing a continuous measure of developmental stage.

---

**Figure S1.**
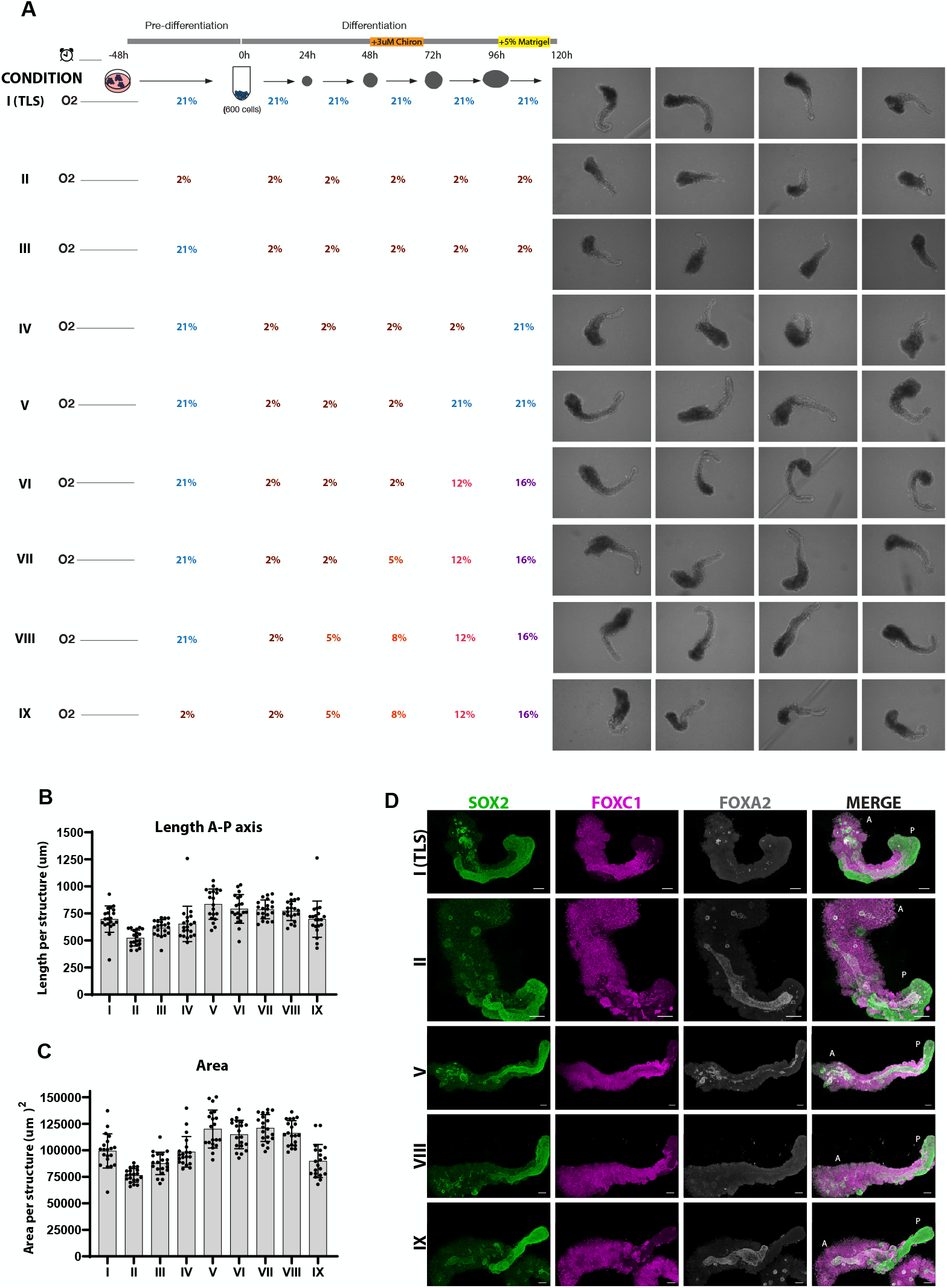
Hypoxia alters TLS morphology and composition. A. Schematic illustration of the experimental pipeline for TLS generation in different O2 setups accompanied by wide-field images. B. Graph of A-P axis length per structure for conditions I-IX. Column height and error bars indicate mean and standard deviation. Each dot represents an individual structure. C. Graph of total area per structure for conditions I-IX. Each dot represents an individual structure. D. IF staining of testing conditions I (conventional TLS), II, V, VIII and IX for neural marker SOX2, somitic marker FOXC1 and endoderm marker FOXA2. A: Anterior, P: Posterior. Scale bars, 100 μm.

**Figure S2.**
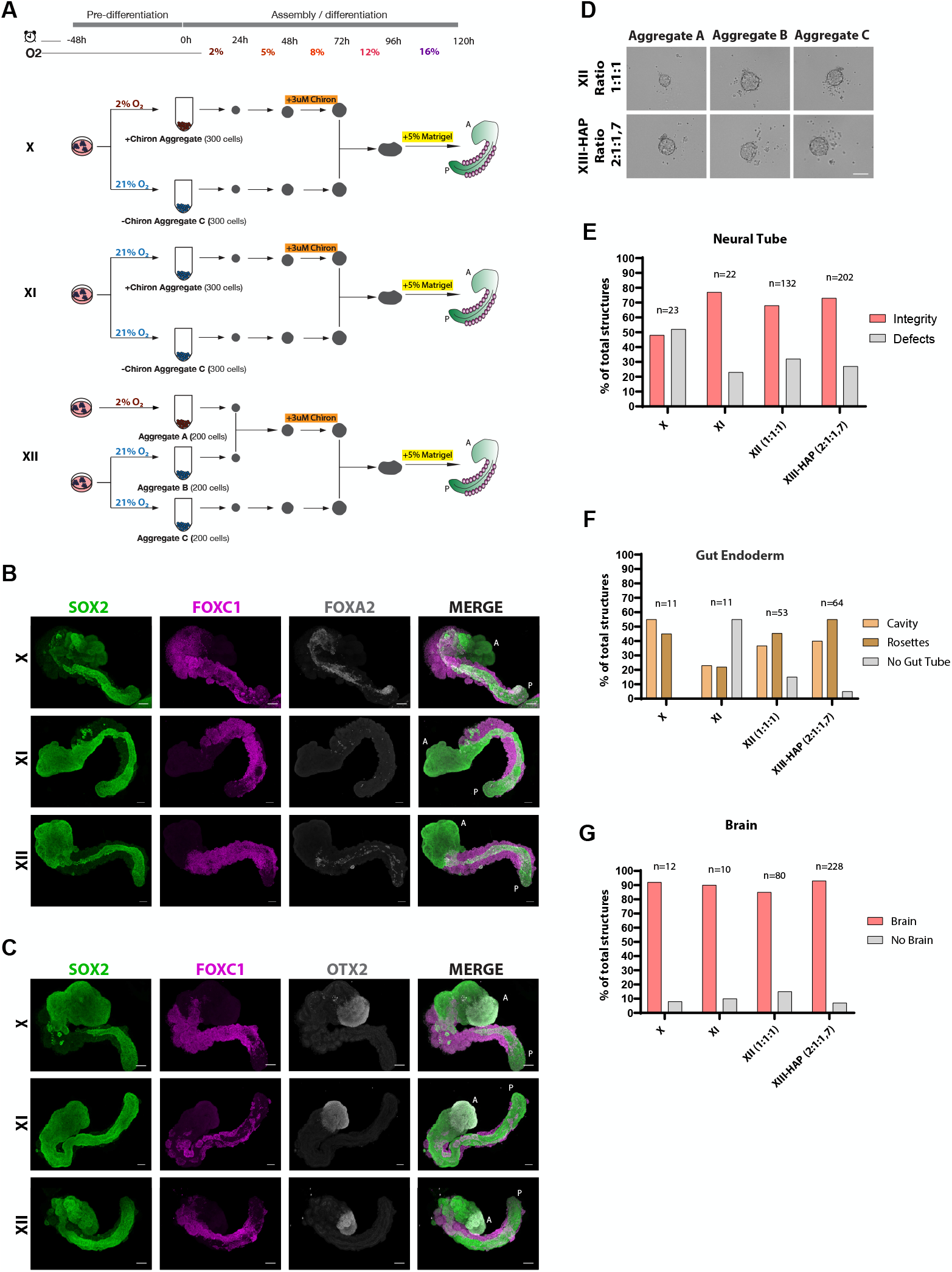
Hypoxic assembloids form anterior neural tissues. A. Schematic illustration of the experimental pipeline for of hypoxic A-P gastruloid optimization steps. B. IF staining for the shown markers. A: Anterior, P: Posterior. Scale bars, 100 μm. C. IF staining for the shown markers. D. Bright-field images of each aggregate of the triple aggregate setup at 24h. Two different ratios were tested in order to achieve equal aggregate size at the 24h time point. E-G. Percentage of total structures showing neural tube integrity, gut endoderm and brain formation in conditions X, XI, XII and XIII (HAP-gastruloids). Column height indicates mean and standard deviation.

**Figure S3.**
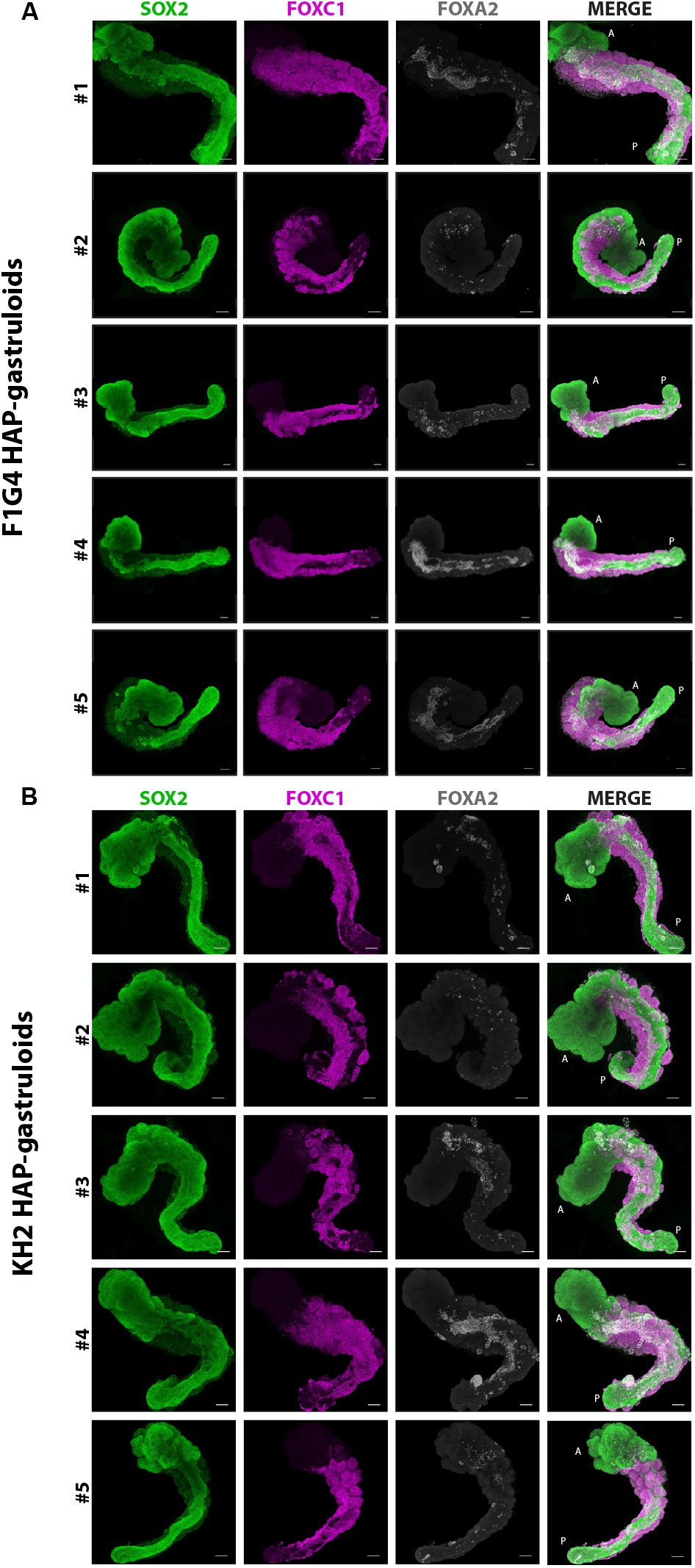
Representative stainings of F1G4 and KH2 HAP gastruloids (different ESC lines) A, B. IF stainings of for the neural marker SOX2, somitic marker FOXC1 and endoderm marker FOXA2. A: Anterior, P: Posterior. Scale bars, 100 μm.

**Figure S4.**
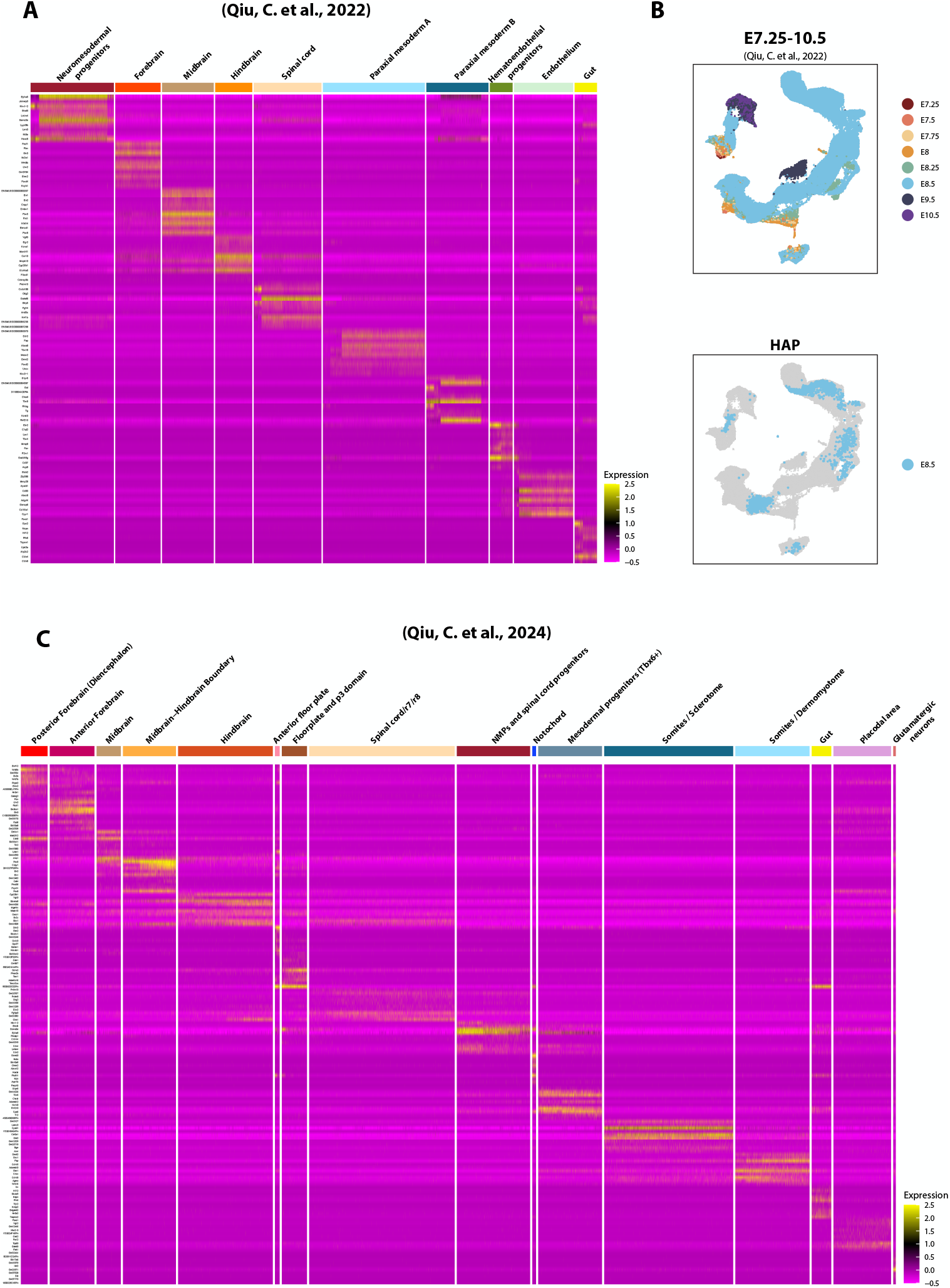
Gene markers for cell type annotation of scRNA-seq data. A. Heatmap showing gene markers for each cell type based on the first reference46. Scaled average expression level is shown. B. Top, UMAP plot of mouse embryo reference scRNA-seq data stages between E7.25 to E10.5 colored by developmental stage. Bottom, UMAP plot of HAP-gastruloid scRNA-seq data projected onto the embryo atlas colored by developmental stage. Only E8.5 stage is detected. C. Heatmap showing gene markers of each cell type based on the second reference47. Scaled average expression level is shown.

**Figure S5.**
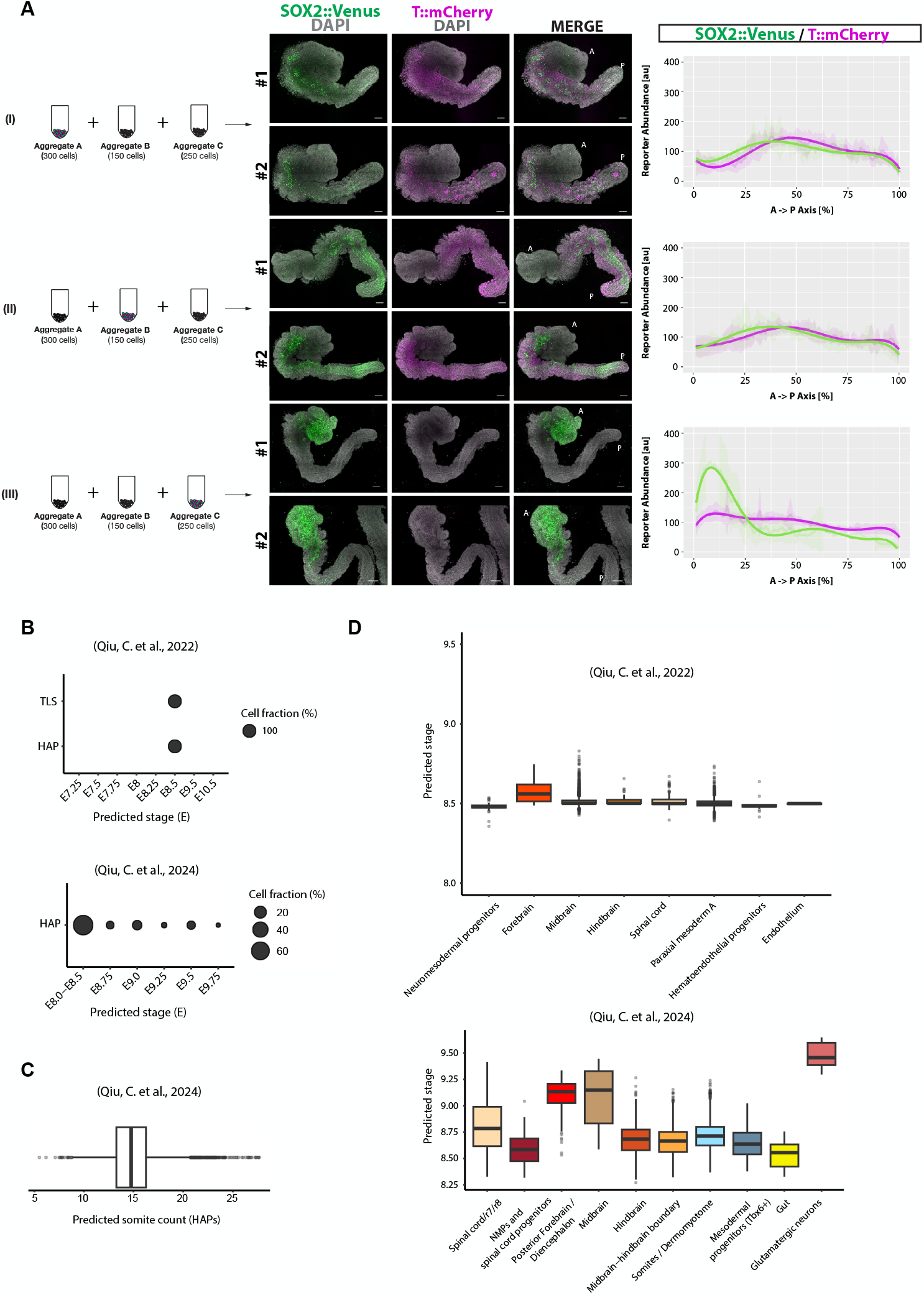
Developmental timing and contribution of tissues derived from each aggregate. A. Contribution of each aggregate in tissue formation in HAP-gastruloids by using a dual reporter cell line for SOX2::Venus and T::mCherry. A: Anterior, P: Posterior. Scale bars, 100 μm. B. Bubble plot showing the predicted developmental progress of HAP-gastruloids and TLSs, based on a label transfer-based staging analysis (see Methods). Bubble size represents the assigned cell fraction in percent. C. Boxplot of predicted somite count in HAP-gastruloids. Line shows median, box borders denote the interquartile range, whiskers show 1.5 times the interquartile range, and dots show outliers. D. Boxplots of predicted developmental stages that each tissue of HAP-gastruloids corresponds to based on label transfer-based staging analysis. Line shows median, box borders denote the interquartile range, whiskers show 1.5 times the interquartile range, and dots show outliers.

**Figure S6.**
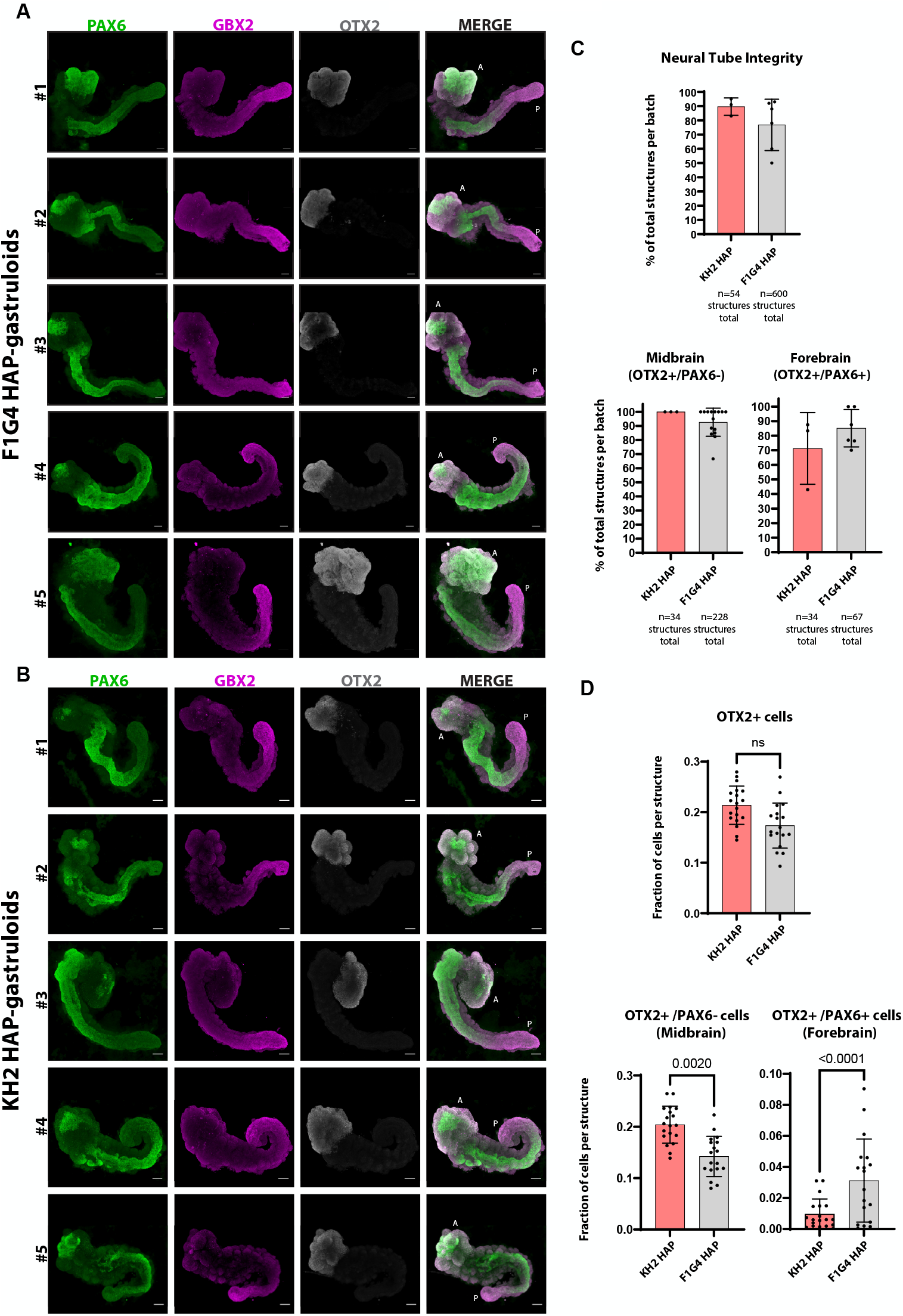
KH2 HAP show appropriate neural patterning along the A-P axis similarly to F1G4 HAP. A, B. IF staining for the shown markers in HAP-gastruloids generated from F1G4 (A) and KH2 (B) ESC lines. A: Anterior, P: Posterior. Scale bars, 100 μm. C. Percentage of total structures per batch showing midbrain and forebrain formation. Column height and error bars indicate mean and standard deviation, respectively. Each dot represents a different batch of structures. D. Fraction of cells per structure expressing the shown markers. Each dot represents an individual structure. Statistical test is unpaired two-tailed t test.

**Figure S7.**
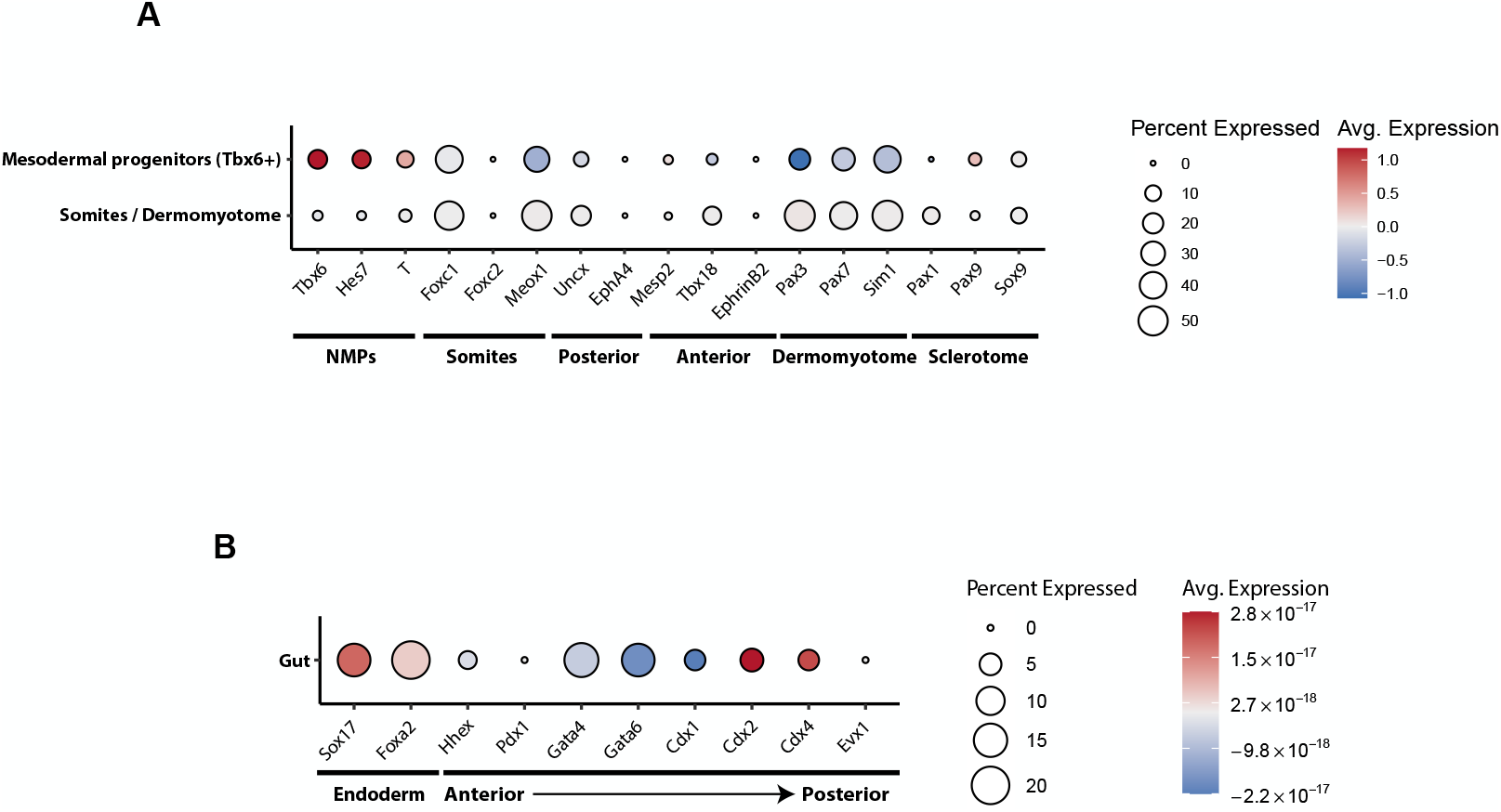
Expression of somite and gut markers in HAP-gastruloids. A. Bubble plot showing the expression of somitic specific marker genes in HAP-gastruloids based on the second reference47. Dot size corresponds to the percentage of cells in the sample expressing the gene, color intensity reflects the scaled average expression level. B. Bubble plot showing the expression of gut specific marker genes in HAP-gastruloids based on the second reference47. Bubble size indicates the percentage of cells expressing the gene, color intensity reflects the scaled average expression level.

**Figure S8.**
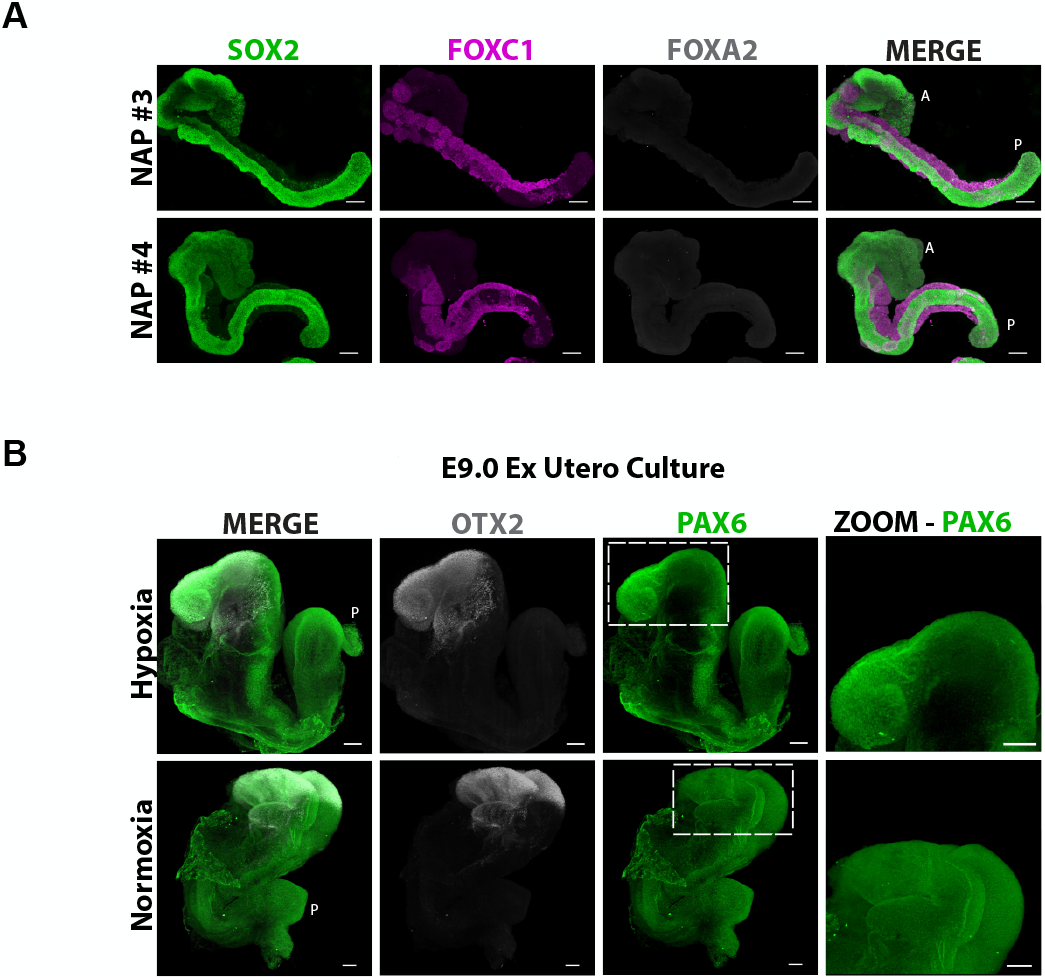
Normoxia impairs gut endoderm and anterior neural fate. A, B. IF staining of NAP-gastruloids (A) and E9.0 ex utero cultured embryos (B) for the neural marker SOX2, somitic marker FOXC1 and endoderm marker FOXA2. A: Anterior, P: Posterior. Scale bars, 100 μm.

**Figure S9.**
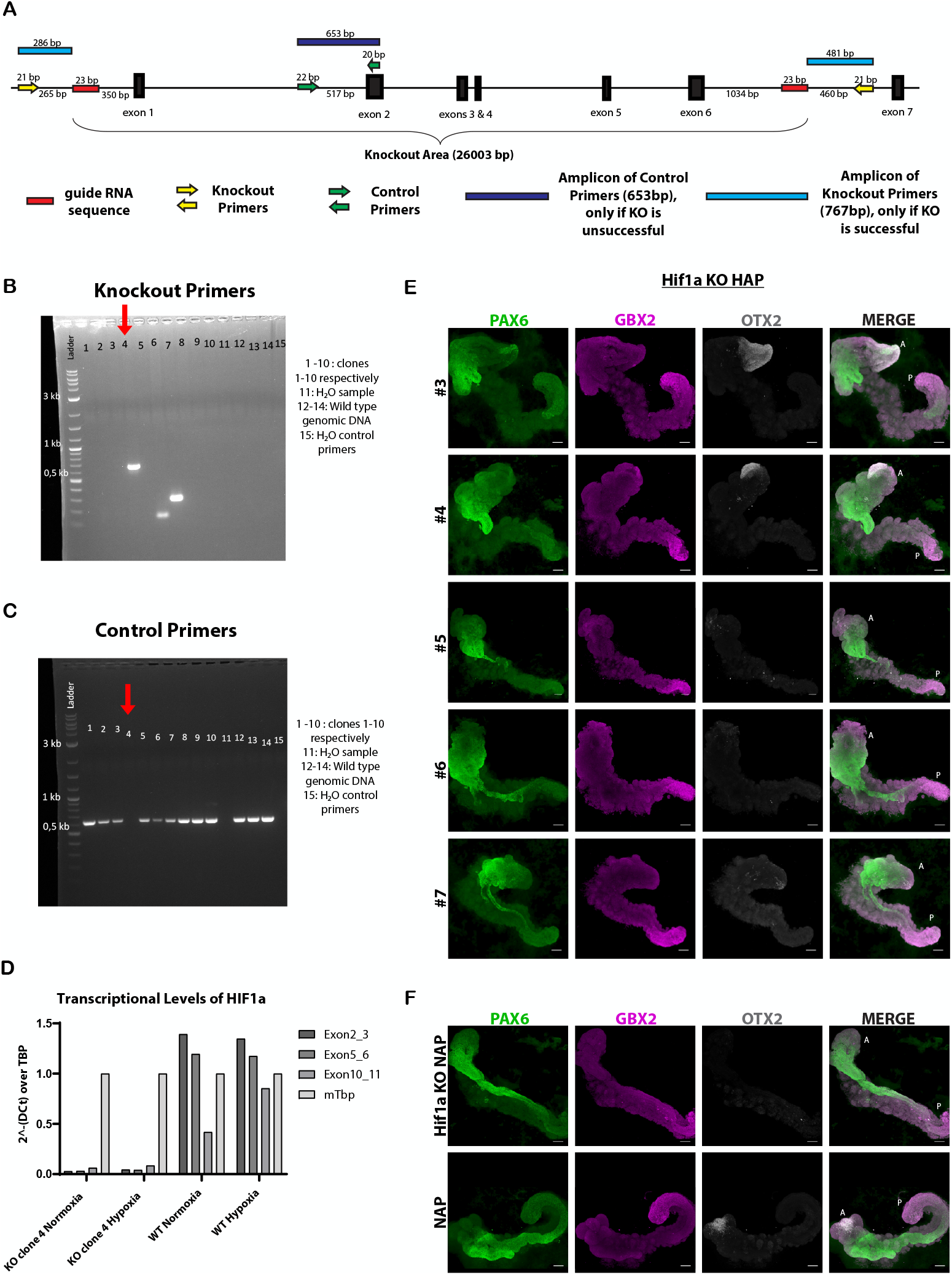
Details of Hif1a KO ESC generation and HAP-gastruloids. A. Schematic illustration of the KO strategy at the Hif1a locus. B, C. Genotyping PCR with knockout (B) and control (C) primers. D. RT-qPCR showing the transcriptional levels of the Hif1a transcript in WT and Hif1a KO cells in hypoxia and normoxia. Three pairs of primers for exons 2+3, 5+6 and 10+11 were used for validation. Mouse TATA-box-binding protein (mTbp) was used as a reference target. The Hif1a gene is post-translationally regulated, therefore transcripts are also detected in normoxia. E. IF staining of additional Hif1a KO HAP-gastruloids for the shown markers. A: Anterior, P: Posterior. Scale bars, 100 μm. F. IF staining of wt and Hif1a KO NAP-gastruloids for the shown markers. A: Anterior, P: Posterior. Scale bars, 100 μm.

**Figure S10.**
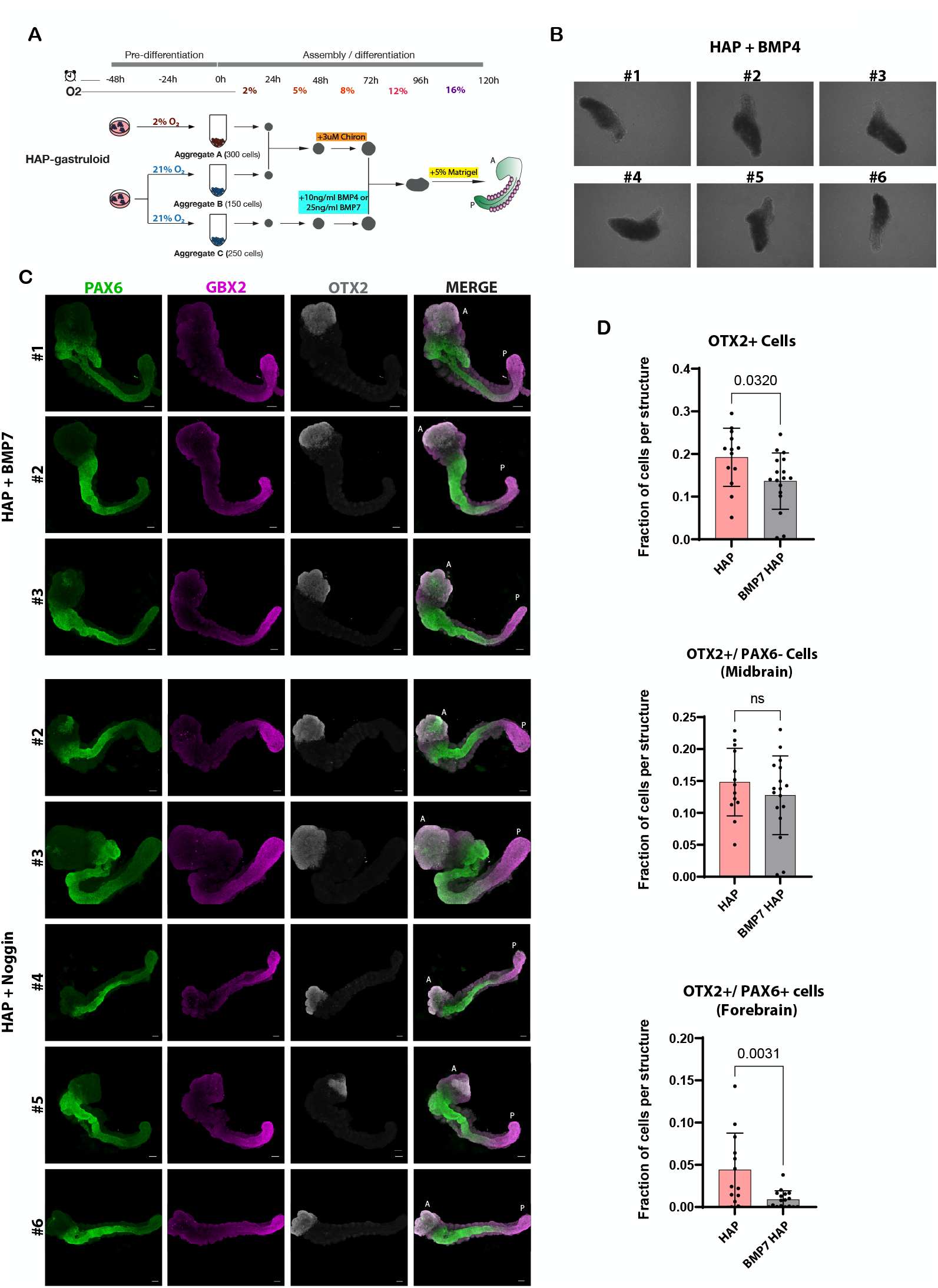
BMP signalling hinders forebrain formation. A. Schematic illustration of the experiment B. Bright-field images of 120h HAP-gastruloids treated with BMP4. C. IF staining of BMP7- or Noggin-treated HAP-gastruloids for the shown markers. A: Anterior, P: Posterior. Scale bars, 100 μm. D. Fraction of cells per structure expressing the shown markers in each condition. Column height and error bars indicate mean and standard deviation. Each dot represents an individual structure. Statistical test is unpaired two-tailed t test.

**Figure S11.**
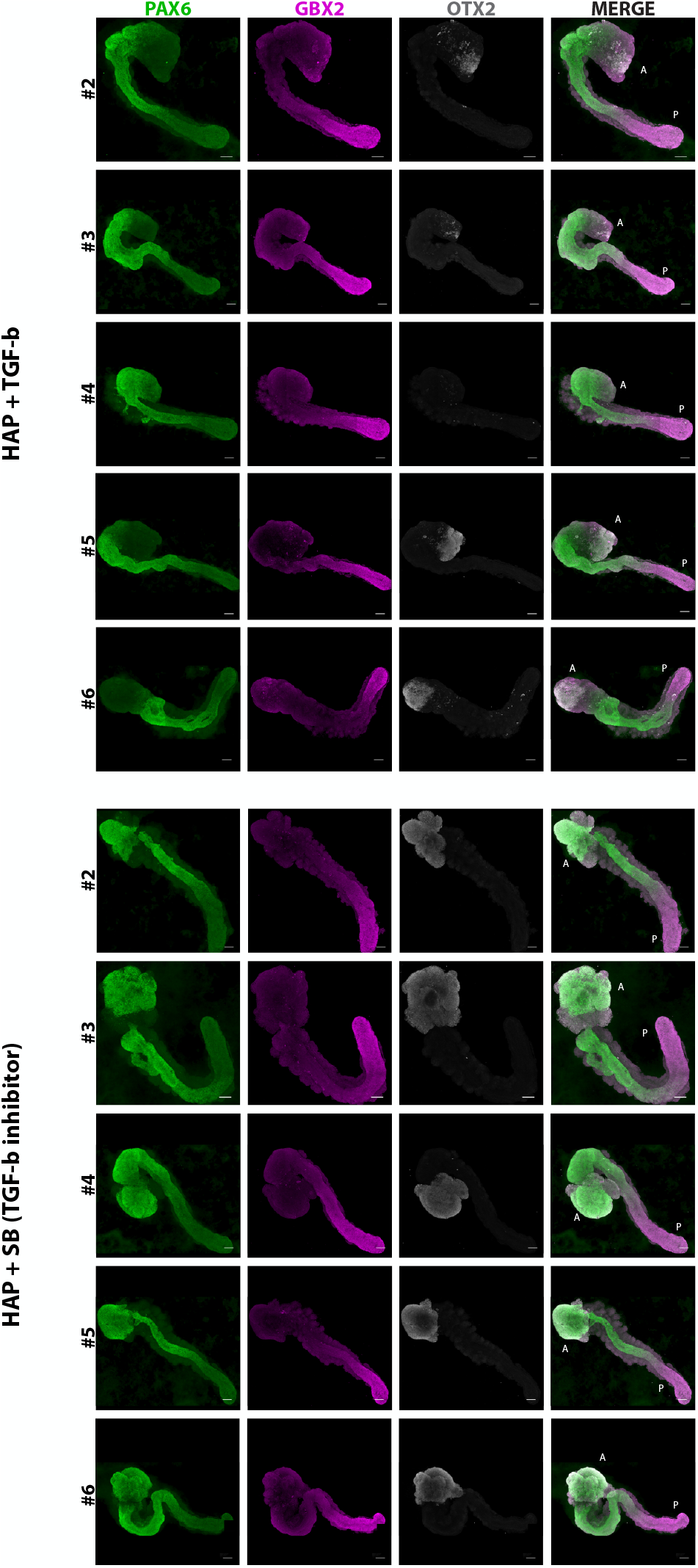
Additional stainings of HAP + TGFβ and HAP + SB. Additional IF stainings of HAP-gastruloids treated with the TGFβ protein or its inhibitor SB for the shown markers.. A: Anterior, P: Posterior. Scale bars, 100 μm.

**Figure S12.**
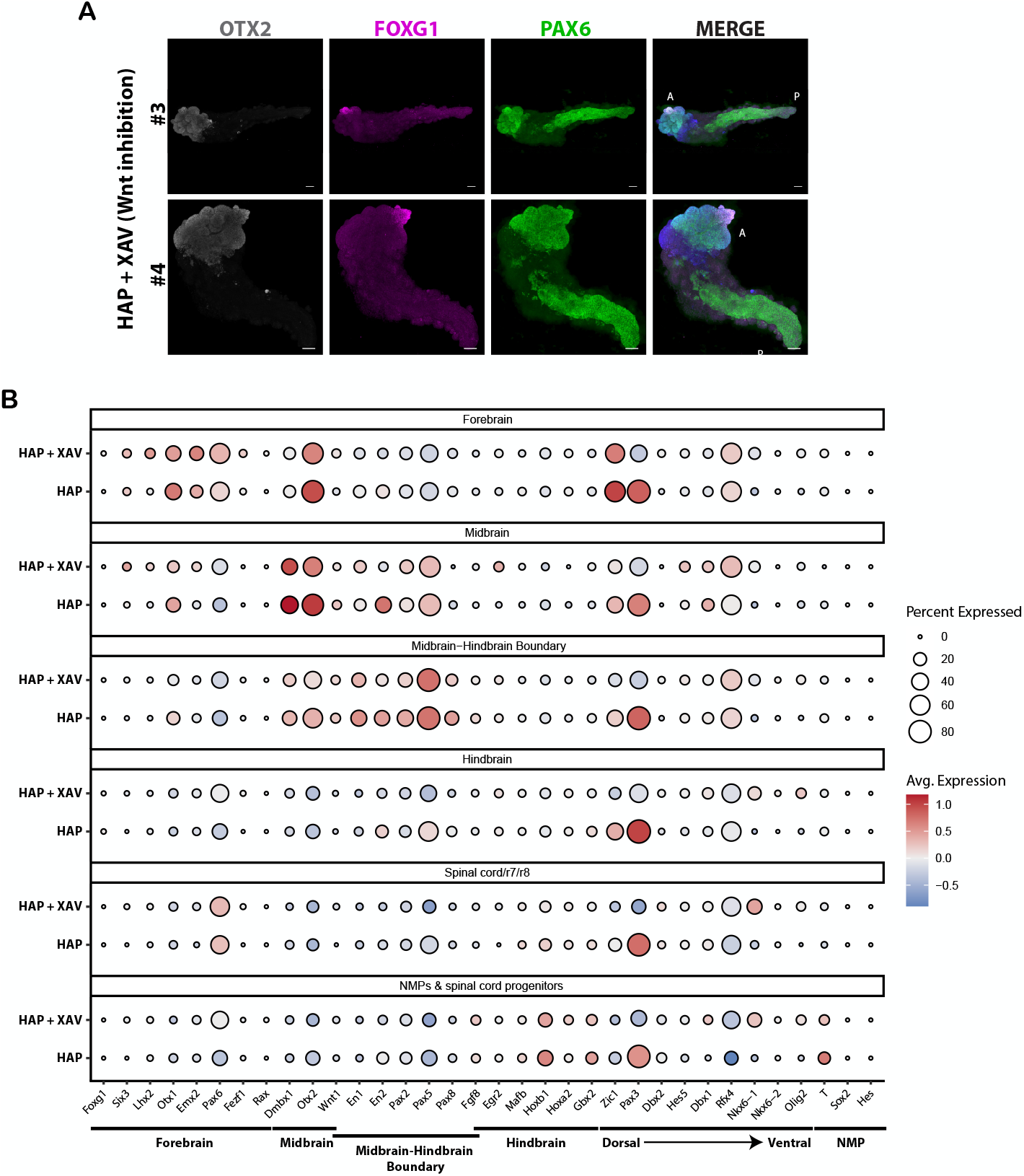
WNT inhibition in HAPs induces anterior forebrain. A. IF staining of HAP-gastruloids treated with the WNT inhibitor XAV for the shown markers. A: Anterior, P: Posterior. Scale bars, 100 μm B. Bubble plot showing the expression of neural cell type specific marker genes47 in HAP-gastruloids treated with XAV compared to untreated ones. Bubble size corresponds to the percentage of cells expressing the gene, color intensity reflects the scaled average expression level.

**Figure S13.**
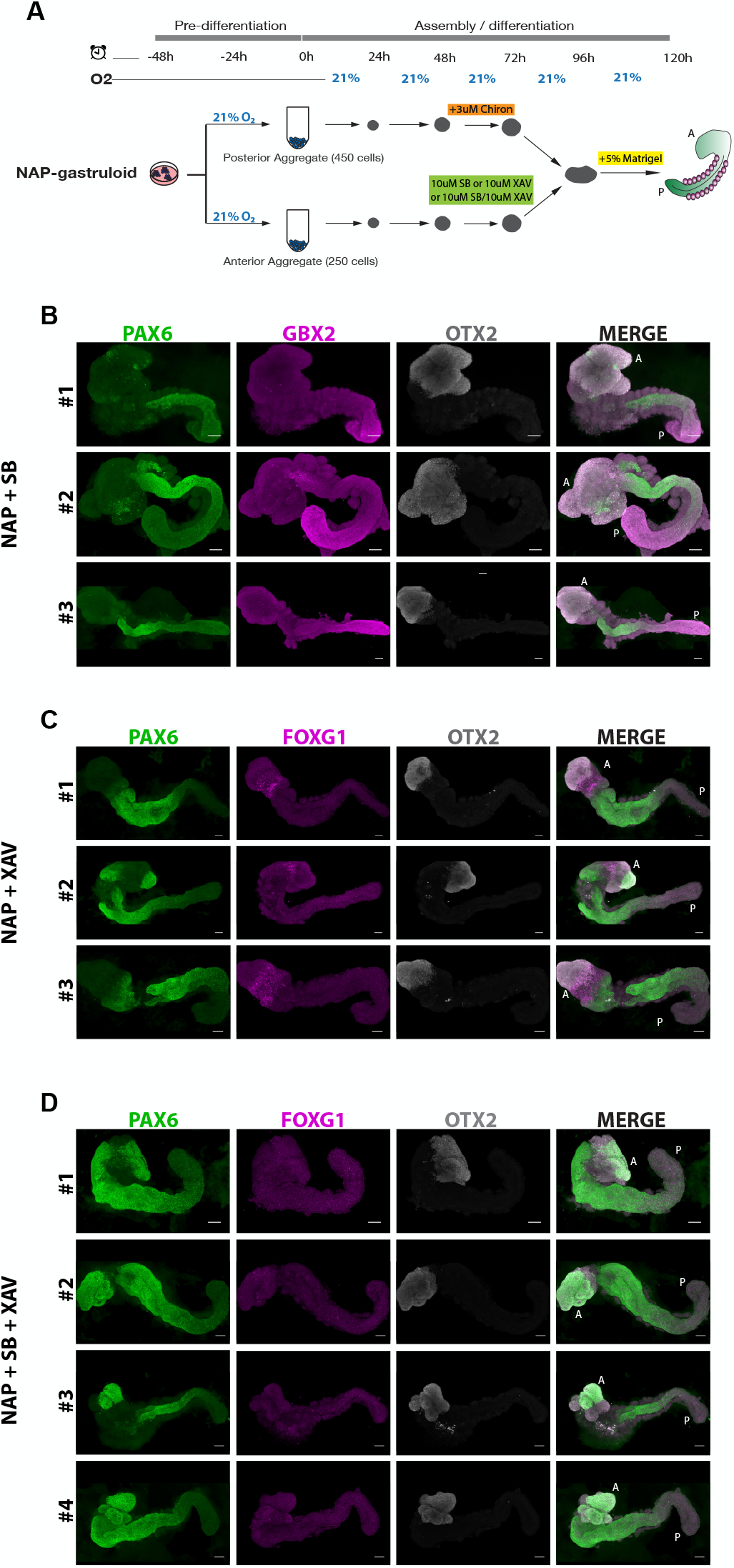
TGFβ and WNT inhibition do not substitute for hypoxia in NAPv. A. Schematic illustration of the experiment. B-D. IF stainings of NAP-gastruloids treated with SB (A), XAV (B), or SB+XAV (C) for the shown markers.. A: Anterior, P: Posterior. Scale bars, 100 μm

